# The alternative oxidase reconfigures the larval mitochondrial electron transport system to accelerate growth and development in *Drosophila melanogaster*

**DOI:** 10.1101/2025.02.20.639223

**Authors:** Geovana S. Garcia, Murilo F. Othonicar, Antonio Thiago P. Campos, Eric A. Kilbourn, Kênia C. Bícego, Johannes Lerchner, Jason M. Tennessen, Marcos T. Oliveira

**Affiliations:** Departamento de Biotecnologia, Faculdade de Ciências Agrárias e Veterinárias de Jaboticabal, Universidade Estadual Paulista “Júlio de Mesquita Filho”, Jaboticabal, SP, Brazil; Departamento de Física, Universidade Federal do Ceará, Fortaleza, CE, Brazil; Department of Biology, Indiana University, Bloomington, IN, United States; Departamento de Morfologia e Fisiologia Animal, Faculdade de Ciências Agrárias e Veterinárias de Jaboticabal, Universidade Estadual Paulista “Júlio de Mesquita Filho”, Jaboticabal, SP, Brazil; Institute of Physical Chemistry, TU Bergakademie Freiberg, Freiberg, Germany

**Keywords:** OXPHOS, respiratory chain, cataplerosis, thermogenesis, supercomplexes

## Abstract

The alternative oxidase (AOX) is naturally present in the mitochondrial electron transfer system (ETS) of many organisms but absent in vertebrates and most insects. AOX oxidizes coenzyme Q and reduces O_2_ in H_2_O, partially replacing the ETS cytochrome c segment and alleviating the oxidative stress caused by ETS overload. As successfully demonstrated in animal models, AOX shows potential in mitigating mitochondrial diseases. However, its non-proton-pumping nature may uncouple mitochondria, leading to excessive heat generation and interference with normal metabolism and physiology. Here we show that AOX from the tunicate *Ciona intestinalis* accelerates development of *Drosophila melanogaster*, elevating larval biomass accumulation (primarily due to increased fat), mobility and food intake, without increasing body heat production. AOX intensifies Leak respiration and lowers oxidative phosphorylation efficiency through functional interactions with the mitochondrial glycerol-3-phosphate dehydrogenase (mGPDH). This is associated with increased complex I (CI)-driven respiration and supercomplex formation, higher cellular NAD+/NADH ratios, and an enhanced flux through the central carbon metabolism. Chemical uncouplers and rotenone confirm the roles of mitochondrial uncoupling and CI in the development of AOX-expressing larvae. Thus, AOX appears to be promoting increased growth by reinforcing the larval proliferative metabolic program via an intricate mechanism that reconfigures the larval ETS.

## Introduction

Investigating mitochondrial metabolism and physiology is crucial for understanding how cells and organisms adjust and adapt to various conditions, such as temperature, nutrition, and oxygen availability (Benej et al., 2024; Cormier et al., 2019; Donnelly et al., 2024; Efeyan et al., 2015; Schou et al., 2017; Scott et al., 2019; Sokolova, 2018). Many of these adjustments occur throughout the organism’s development, so gaining insights into these processes and the mechanisms of metabolic resilience can provide valuable biological knowledge that can even contribute to our understanding of the molecular basis of human disorders (Drummond-Barbosa and Tennessen, 2020).

While mitochondria have a myriad of roles in cellular metabolism and beyond, they are traditionally famous for housing the bulk synthesis of ATP by means of oxidative phosphorylation (OXPHOS). This process integrates various cellular metabolic pathways with an electron transfer system (ETS) in the inner mitochondrial membrane (IMM), allowing the transduction of the chemical energy stored in nutrients into utilizable forms of energy. The ETS is composed of different dehydrogenases, the most studied being respiratory complexes I (CI) and II (CII), which oxidize reduced metabolites, converging the electrons to IMM’s coenzyme Q molecules (CoQ) (Banerjee et al., 2021; Larosa and Remacle, 2018; Nolfi-Donegan et al., 2020). A series of redox reactions through complexes III (CIII) and IV (CIV) leads to CoQ reoxidation and the reduction of molecular oxygen, which is coupled to the formation of an electrochemical gradient across the IMM by proton (and sodium) pumping through CI, III and IV (Larosa and Remacle, 2018; Mitchell, 1961; Nolfi-Donegan et al., 2020; Pablo Hernansanz-Agustín et al., 2024). These three complexes are often found physically interacting in supramolecular organizations generally termed as supercomplexes (SC), which may stabilize each individual complex and enhance electron transfer and ion pumping (Lapuente-Brun et al., 2013; Milenkovic et al., 2017; Schagger, 2000). ATP synthase then uses the potential energy (ΔΨm) of the electrochemical gradient to produce ATP, but notably, this energy can also be released as heat, as protons return to the mitochondrial matrix independently of ATP synthase (Divakaruni and Brand, 2011; Mitchell, 1961; Nicholls, 2021; Zhao et al., 2019).

The balance of proton gradient dissipation between ATP synthesis and heat production can be regulated. In the mammalian brown adipose tissue, low levels of ATP synthase and high levels of the uncoupling protein UCP1 result in less efficient OXPHOS and in non-shivering thermogenesis, protecting small animals, including neonatal humans, from hypothermia (Cannon and Nedergaard, 2004; Matthias et al., 2000; Nedergaard et al., 2001; Porter, 2017; Porter et al., 2016). Non-shivering thermogenesis caused at least in part by mitochondrial uncoupling is also observed in the skeletal muscle of the tegu lizard during its high-demand reproductive season (Hervas et al., 2024). In this case, the uncoupling is due to the adenine nucleotide translocase (ANT) activity, and not UCP. In plants, alternative enzymes (AEs) in the IMM act by uncoupling electron transfer to molecular oxygen from proton pumping, thus compromising the formation of the electrochemical gradient used by the ATP synthase (Borecký and Vercesi, 2005; Juszczuk and Rychter, 2003; Rogov et al., 2014; Watling et al., 2006). Their “alternative” status comes mostly from the fact that they can either transfer electrons from the soluble electron carrier NADH to CoQ (a reaction catalyzed by the alternative NADH dehydrogenase, which is equivalent to that catalyzed by CI) (Matus-Ortega et al., 2011; McDonald and Gospodaryov, 2019), or transfer electrons from CoQ directly to molecular oxygen (similarly to the concerted actions of CIII and IV, but instead catalyzed by a single enzyme, the alternative oxidase AOX) (Juszczuk and Rychter, 2003; McDonald and Gospodaryov, 2019). As a result, the energy of nutrient oxidation is dissipated as heat, serving as a thermogenic mechanism that permits some species to flower under extremely low temperatures or to volatilize pollinator-attracting odors (Inaba et al., 2019; Ito et al., 2004; Onda et al., 2008; Watling et al., 2006).

Interestingly, AEs are present in most plant, fungi and animal groups, but are naturally absent in vertebrates and insects (with the exception of sciarid dipterans) (McDonald and Gospodaryov, 2019; Weaver and McDonald, 2023). They have, however, been xenotopically expressed in several biological models as an approach to explore their potential use as therapeutic tools for human mitochondrial dysfunction (El-Khoury et al., 2016, 2013; Fernandez-Ayala et al., 2009; Giordano et al., 2019; Hakkaart et al., 2006; Humphrey et al., 2012; Jacobs et al., 2023; Kemppainen et al., 2014; Rajendran et al., 2019; Saari et al., 2019a; Sanz et al., 2010; Szibor et al., 2020a, 2020b), which may cause classical mitochondrial diseases or contribute to the outcome of complex diseases, such as cancer, neurodegeneration, obesity, etc. (Bhatti et al., 2017; Herst et al., 2017; Murphy and Hartley, 2018; Weinberg and Chandel, 2015). In a mitochondrial system like in humans, where CoQ reduction by NADH and CoQ reoxidation by molecular oxygen depend solely on the activities of CI and CIII/IV, respectively, inhibition of these respiratory complexes—whether genetic, physiological, or chemical—compromises the ETS and may result in the production of toxic levels of reactive oxygen species (ROS) (Bhatti et al., 2017; Brzezinski et al., 2021; Chenna et al., 2022; Fiedorczuk and Sazanov, 2018; Ghezzi and Zeviani, 2018; Hadrava Vanova et al., 2020; Szczepanowska et al., 2020). The expression of AEs in various diseases and pathological conditions could then maintain electron flow to molecular oxygen, bypassing eventual unproductive endogenous CI, III and/or IV.

The AOX of the tunicate *Ciona intestinalis* is the AE that has been best studied in heterologous systems, with numerous results indicating the positive outcomes of its expression in different models of mitochondrial dysfunction (Andjeiković et al., 2016; Andjelković et al., 2018; El-Khoury et al., 2016, 2013; Fernandez-Ayala et al., 2009; Giordano et al., 2019; Hakkaart et al., 2006; Humphrey et al., 2012; Jacobs et al., 2023; Kemppainen et al., 2014; Mills et al., 2016; Rajendran et al., 2019; Szibor et al., 2020b). At least part of these outcomes may be attributable to the enzyme’s pro-thermogenic properties, since its expression does mitigate the effects of cold stress on the model organism *Drosophila melanogaster*, contributing to a fast growth under low temperatures (Saari et al., 2019a), and mitochondria-generated heat might be at the core of several important biological processes (Chrétien et al., 2018; Macherel et al., 2021; Terzioglu et al., 2023).

However, there have been a few key reports showing severe negative impacts of AOX expression in both *Drosophila* and mice (Dogan et al., 2018; Saari et al., 2019a, 2019b, 2017). In the former, these harmful effects can be observed in situations with functionally intact ETS complexes, i.e., without classical mitochondrial dysfunction.

For example, mature sperm production by adult males is seriously affected by AOX expression in the testis in a dose-dependent manner, a problem that only becomes apparent when these males are challenged in sperm competition assays (Saari et al., 2017). In addition, fly development upon AOX expression relies on a nutrient-rich diet, without which most individuals die at the pupa stage (Saari et al., 2019a, 2019b), preceded by defects in muscle function at the larval stage (Couto-Lima et al., under review). The detrimental effects of AOX on *Drosophila* development might be, at least in part, consequences of a “metabolic interference” between fly supercomplexes and AOX, since this enzyme appears not to encounter such supramolecular structures in its “natural habitat”, the *C. intestinalis* IMM (Othonicar et al., 2024). Therefore, how *C. intestinalis* AOX and other AEs influence mitochondrial metabolism and whole organism physiology/phenotype may vary significantly across cell/tissue types, developmental stages and environmental conditions, highlighting our need to better understand the metabolic contexts within which these enzymes can be deployed as treatments, and which may reflect their natural IMM environments.

Here, we explore the mechanisms by which *C. intestinalis* AOX interferes with the *Drosophila* ETS, triggering metabolic resilience during larval growth and development, particularly under stressful cold temperatures. Based on its non-proton pumping nature and its roles in plant thermogenesis, we hypothesized that this effect of AOX expression in flies is due to increased thermogenesis via mild mitochondrial uncoupling. To test this, we quantified larval and pupal body temperatures and phenotyped several developmental parameters, combining these data with larval metabolic and mitochondrial physiology data. We specifically analyzed two distinct ETS pathways: (i) the one initiated by CI with oxidation of mitochondrial matrix-generated NADH, which can operate in supercomplexes with CIII and IV (Milenkovic et al., 2017; Schagger, 2000); and the pathway initiated by the mitochondrial glycerol-3-phosphate dehydrogenase (mGPDH), a FAD-linked enzyme facing the mitochondrial intermembrane space that reduces CoQ by oxidizing glycerol-3-phosphate (G3P), a product of intermediary metabolism bridging glycolysis and lipid metabolism (Mráček et al., 2013). Our results reveal an intricate mechanism by which AOX preferentially uncouples mitochondria via mGPDH-driven respiration while simultaneously increasing CI-driven respiration via increased supercomplex formation. In turn, AOX expression stimulates cellular NADH reoxidation and mitochondrial cataplerosis, thus enhancing the larval metabolic growth program.

## Results

### AOX contributes to increasing larval biomass accumulation and growth

We previously demonstrated that AOX both increases viability and decreases the egg-to-pupa developmental time in *Drosophila* cultured at low temperatures (Saari et al., 2019a). To determine if these effects could be attributable to increased thermogenesis, we examined L3 larval and prepupal body temperatures in AOX-expressing flies and in a genetically matched control line. We used the previously described fly line *3XtubAOX* (Kemppainen et al., 2014; Saari et al., 2017), which expresses *C. intestinalis* AOX constitutively under the control of the *α-tubulin* promoter, and the background line *w^1118^* as control. We first confirmed that the expression of the *AOX* transgenes did not change due to temperature (Supplemental Figure S1A) and demonstrated that the faster developmental time of *3XtubAOX* larvae was not caused by the extra copies of the *white* gene in this line, used as markers for the transgenesis (Supplemental Figure S1B).

As a first approach to analyzing thermogenesis, we utilized a specialized thermopile chip calorimeter designed for small biological samples (Lerchner et al., 2024), and measured real-time larval and prepupal heat production at the constant temperatures of 25 and 13 °C. We observed no significant differences in the average heat production between control and *3XtubAOX* larvae and prepupae (Figure 1B).

**Figure 1.**
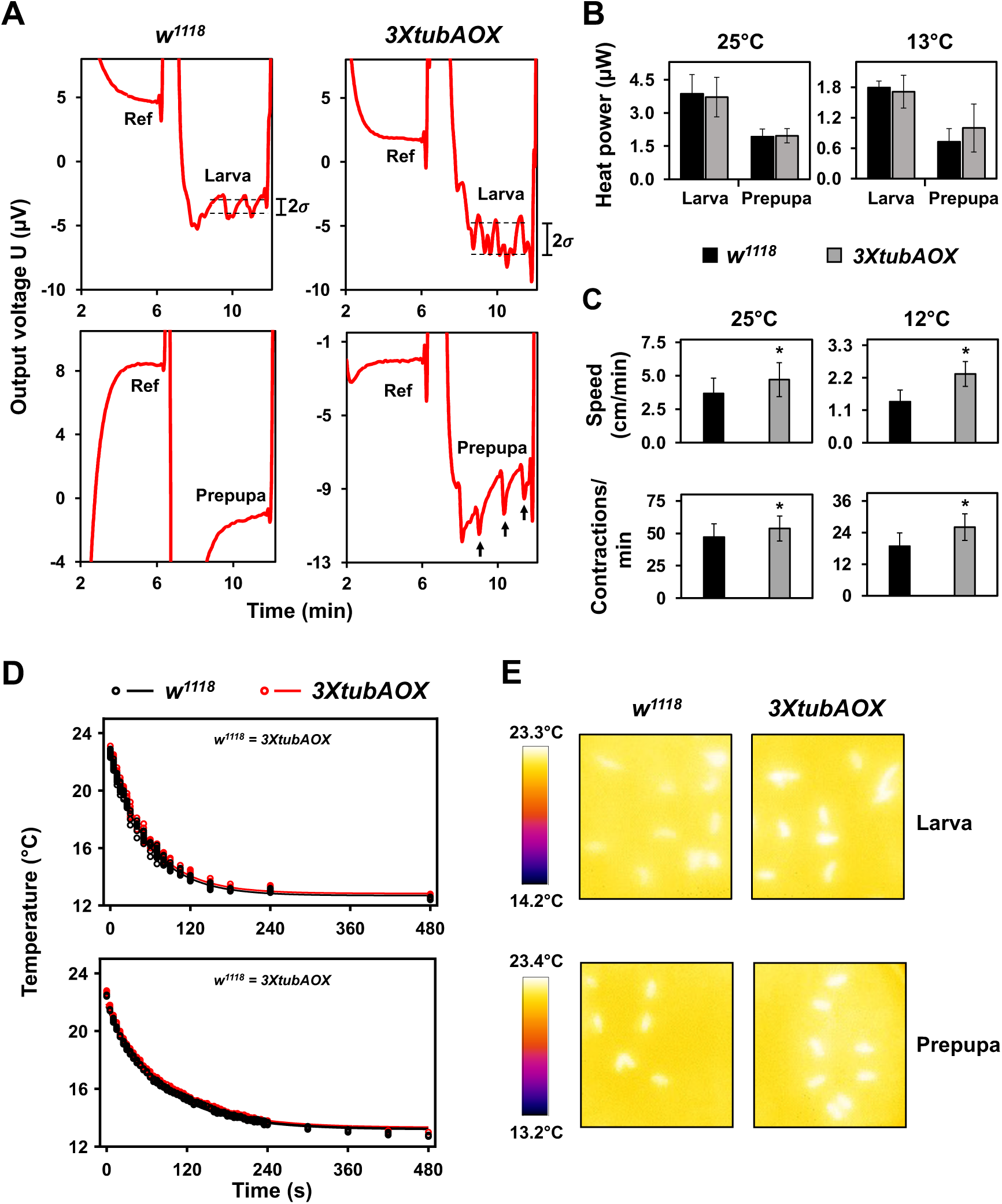
Larval and prepupal heat production and body temperatures. (A) Representative output of real-time voltage changes in thermopiles of the small-sample chip calorimeter described by (Lerchner et al., 2024). Top graphs, signals obtained for *Drosophila* larvae, showcasing the noise attributed to movement (2σ). Quantitation of 2σ is shown in Supplementary Table S1. Bottom graphs, signals for prepupae; note that while control prepupae exhibit a stable heat output signal, *3XtubAOX* prepupae show frequent heat bursts (black arrows), which are not attributed to mobility. Quantitation of heat burst events is also shown in Supplementary Table S1. Ref, a stable signal measured with an inert sample in a segment of the thermopile, used as reference. (B) Average heat production of larvae and prepupae at 25 or 13 °C, calculated as the voltage difference related to the Ref segment. Data represent means +/-standard deviation of 6-14 individuals. (C) Distance crawled and body wall contractions per min of control and *3XtubAOX* larvae culture at 25 and 12 °C. Data represent means +/-standard deviation of 10-15 technical replicates from 10 biological replicates for each genotype. (D) Infrared thermographic measurements of changes in whole-body temperature over time in control (*w^1118^*) and AOX (*3xtubAOX*) larvae (top panel) and prepupae (bottom panel). Animals cultured at 25 °C were transferred to 12 °C and filmed using a thermographic camera, as described in the Materials and Methods. Each data point represents the body temperature of an individual animal from a biological replicate, and the curves represent the 1^st^ order exponential fitting curve of temperature decay. (E) Representative infrared thermography images show control flies and AOX flies immediately after placement at 12 °C (time 0).

However, *3XtubAOX* larvae showed a twofold increase in the standard deviation of the signal noise (Figure 1A, indicated by 2σ, and Supplemental Table S1). This elevated noise in the output of the calorimetric measurements is attributed to the intense mobile activity of the *3XtubAOX* larvae, which is consistent with our data showing that AOX elevates larval crawling speed and peristaltic body contractions at 25 and 12 °C (Figure 1C, and (Garcia et al., 2021)). Intriguingly, we also observed that *3XtubAOX* prepupae frequently display short heat bursts occurring at ∼1 min intervals (Figure 1A, black arrows), with these events two-threefold more likely to occur in *3XtubAOX* prepupae than in controls (Supplemental Table S1). To our knowledge, these heat bursts have not been described previously.

We also measured via infrared thermography body temperature of flies originally cultured at 25 °C that were acutely exposed to 12 °C (Figures 1D and E). The temperature decay as a function of time for both control and AOX flies follows 1^st^ order exponential curves (Figures 1D), suggesting similar rates of heat dissipation and body temperatures. Temperature differences, if any, would be mainly due to different heat capacities, as the cooling of the Petri dishes forces a heat flow from the larvae/pupae to the environment, leading to excess temperature because of a non-negligible thermal resistance between the larvae/pupae and the dishes. To gain further inferences of body heat capacity and heat exchange rates, we first calculated larval mass, water, fat and protein content. We observed that *3XtubAOX* larvae have increased wet and dry masses, especially at 12 °C (∼19% more in wet and ∼33% more in dry mass than *w^1118^*) (Figure 2A). Note that the extra copies of the *white* gene in *3XtubAOX* larvae also did not interfere with body masses (Supplemental Figure S1C). Body length and width of AOX larvae were also larger, regardless of temperature (Figure 2B and C). The increase in biomass in AOX larvae can be primarily attributed to an increase of 40 and 60% of total fat mass at 25 and 12 °C, respectively (Figure 2A). This represented a consistent increase of ∼20% in relative fat mass, irrespective of temperature (Figure 2B). The fat body of *3XtubAOX* larvae appears to accumulate more triglycerides per cell, with a slight increment in the size of lipid droplets (Figures 2E-G). Lean mass and relative protein mass did not significantly differ between fly lines, except for a ∼20% elevation in lean mass in the AOX larvae at 12 °C (Figures 2A).

**Figure 2.**
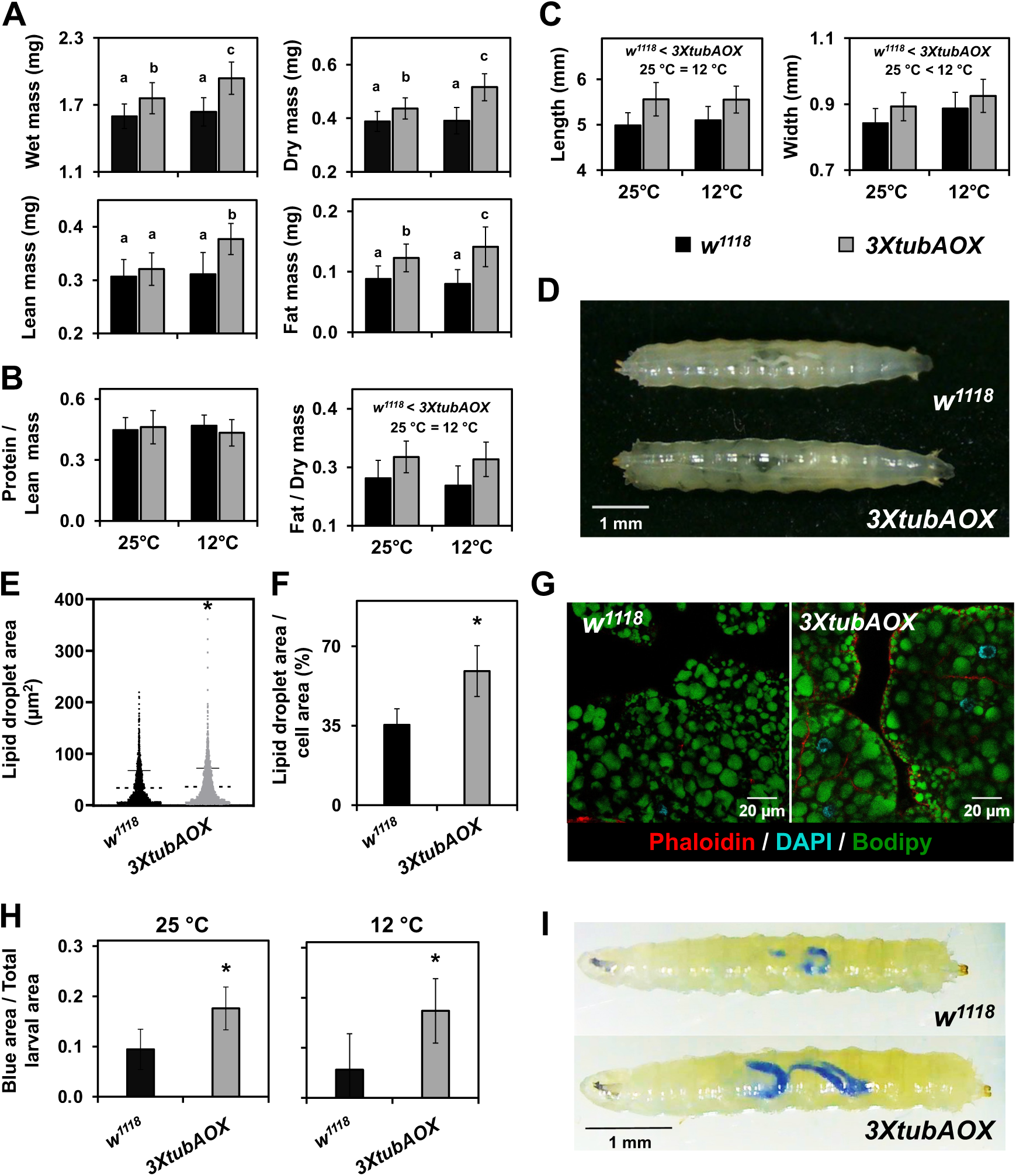
**Larval biomass, size and food intake are increased by AOX. (**A, B) Total and relative masses, and (C) length and width of larvae, were measured as described in the Materials and Methods. Data are presented as means ± standard deviation (SD) from three biological replicates, each comprising at least 24 groups of 10 larvae. Significant differences are denoted as follows: “*w^1118^* < *3XtubAOX*”, differences between genotypes, and “25 °C < 12°C”, differences between temperatures. Letters ‘a–c’ indicate distinct statistical classes based on significant genotype-temperature interactions, determined by two-way ANOVA followed by Tukey’s post hoc tests (p < 0.05). (D) Representative light microscopy images of two wandering female L3 larvae cultured at 12°C. (E) Quantification of lipid droplets in fat body cells of wandering L3 larvae cultured at 25 °C, based on three biological replicates. Error bars represent ± SD of the mean; p < 0.05, based on unpaired t-test with Welch’s correction. (F) Lipid droplet area relative to total cell area. Data are presented as means ± SD from 4–5 independent samples. * indicates p < 0.05 according to Student’s t-test. (G) Representative confocal microscopy images of fat body cells showing lipid droplets. (H) Food intake, quantified as the percentage of the larval gut area stained with blue dye relative to the total L3 larval area (see Materials and Methods for details). Data represent means ± SD from 13-19 individuals sampled across two biological replicates. * denotes significant differences (p < 0.05), based on Student’s t-tests. (I) Representative light microscopy images of L3 larvae after ingesting dyed food.

Heat capacity of *w^1118^* and *3XtubAOX* larvae was, respectively, 5.68 and 6.37 mJ/K. In addition, using the cooling rate of 0.26 K/s (derived from the data in Figure 1C), we calculated the heat exchange rates of 1.48 mJ/s for *w^1118^* and 1.66 mJ/s for *3XtubAOX*. These differences are considerably higher than any difference in body temperature and heat production, and are directly proportional to the increased larval mass, water and fat content in AOX flies. We then evaluated whether the enzyme influences food intake and observed that the *3XtubAOX* larvae ingest two to threefold as much food in our assays as the control larvae (Figure 2H and I). In summary, our data does not support a model in which AOX expression induces elevated thermogenesis (at least not detectable at this point) via mitochondrial uncoupling. However, it is noticeable that AOX increases larval heat capacity and heat exchange rates due to increased body mass and enhanced growth, which is sustained by elevated larval food consumption.

### AOX reconfigures the larval mitochondrial respiratory system

We next examined if AOX could alter larval mitochondrial oxygen consumption rates (OCR) and respiratory control ratios linked to CI and mGPDH substrate oxidation. These are the two most important IMM dehydrogenases of the *Drosophila* larvae, accounting for the large majority of ETS activity (data not shown). In the OXPHOS state (when functional ATP synthase facilitates the proton gradient dissipation) at 25 °C, simultaneous stimulation of CI and mGPDH (CI + mGPDH) and stimulation of CI only resulted in comparable OCR between *3XtubAOX* and control larvae (Figure 3A). On the other hand, mGPDH-driven OCR was 30% decreased in *3XtubAOX* larvae, consistent with our previously published data (Othonicar et al., 2024). At 12 °C, the same drop of 30% in mGPDH-driven OCR in *3XtubAOX* larvae was observed, whereas OCR driven by CI + mGPDH or by CI only increased significantly in the presence of AOX (Figure 3A).

**Figure 3.**
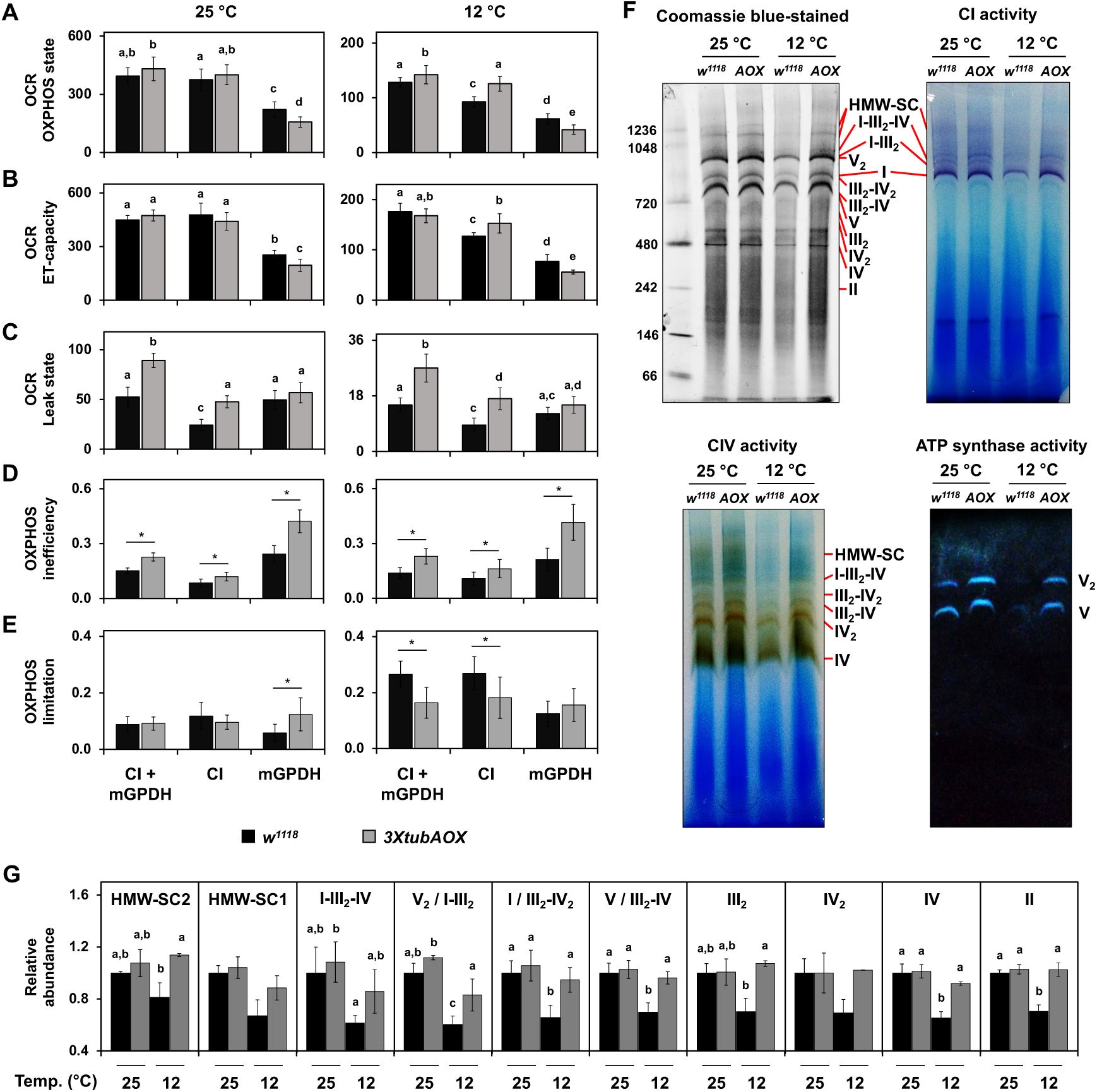
Reconfiguration of the larval mitochondrial respiratory system in the presence of AOX. Oxygen consumption rates (OCR, pmol / s * mg total protein) of wandering L3 larvae at 25 and 12 °C driven by Complex I (CI) and mitochondrial glycerol-3-phosphate dehydrogenase (mGPDH) were measured, respectively, after addition of pyruvate/malate and glycerol-3-phosphate/rotenone. The OXPHOS (A) and ET-capacity (B) states were achieved, respectively, after addition of ADP and a titration of the uncoupler carbonyl cyanide m-chlorophenylhydrazone. The Leak state (C) was obtained in separate assays following the addition of the ATP synthase inhibitor oligomycin. Data points represent the mean +/-standard deviation of 6 biological replicates. Letters ‘a–c’ indicate distinct statistical classes based on significant genotype X pathway(s) interactions, determined by two-way ANOVA followed by Tukey’s *post hoc* tests (p < 0.05). (D) OXPHOS inefficiency, calculated as the ratio Leak OCR / OXPHOS OCR, represents the OXPHOS coupling-control ratio (Gnaiger, 2020) and estimates the proportion of respiration not used in ATP production. (E) OXPHOS limitation, calculated as 1 – (OXPHOS OCR / ET-capacity OCR), expresses the limitation imposed on the electron transport system by the ADP phosphorylation system. Data points represent means +/-standard deviation of at least 4 biological replicates. * indicates significant differences (p < 0.05) according to Student’s t-tests. (F) Representative blue-native polyacrylamide gels showing the separation of mitochondrial complexes and supercomplexes from wandering L3 larvae (*w^1118^*, control line; AOX, *3XtubAOX* line) grown at 25 °C and 12 °C. Mitochondria were solubilized with digitonin, and gels were either stained with Coomassie Blue or analyzed by in-gel activity. These analyses, combined with literature data, confirmed the positions of the indicated complexes. Labeled bands represent individual complexes (I-V), dimers (denoted by a subscripted “2”), or supercomplex structures. HMW-SC refers to high molecular weight supercomplexes. (G) Quantification of Coomassie-stained gels for the indicated complexes, dimers, or supercomplexes was performed via band densitometry. The density of each band was normalized to the density of the control at 25 °C within the corresponding lane. Data are presented as means ± standard deviation from two-three biological replicates. Letters ‘a–c’ indicate distinct statistical classes based on significant genotype X temperature interactions, determined by two-way ANOVA followed by Tukey’s *post hoc* tests (p < 0.05).

We also investigated the active participation of AOX in the OXPHOS respiratory state through its inhibition with propyl-gallate (PG) (Supplemental Figure S2A). At 25 °C, respiration driven by CI + mGPDH or by CI only in *3XtubAOX* larvae was significantly more sensitive to PG, while respiration via mGPDH was not affected. At 12 °C, addition of PG caused significant drops in the OCR initiated via CI or mGPDH, independently or concomitantly, in *3XtubAOX* larvae. In parallel experiments, we verified antimycin A (AA)-resistant respiration, to show the maximum OCR maintained by AOX in the absence of CIII (Supplemental Figure S2B). While AA abolishes all mitochondrial respiration in controls, in *3XtubAOX* larvae 30-40% of OXPHOS remains both at 25 and 12 °C, regardless of whether it occurs via CI and/or mGPDH. Our results indicate that AOX reconfigures and actively participates in the larval mitochondrial respiration.

We also assessed two other respiratory states with opposite effects on ΔΨm: ET-capacity and Leak, during which ΔΨm is respectively minimal and maximal. The ET-capacity state, achieved by titration of the protonophore carbonyl cyanide m-chlorophenyl hydrazone (CCCP) after the OXPHOS state, shows that maximal electron transfer initiated by CI and/or mGPDH in *3XtubAOX* larvae has very similar pattern as seen for the OXPHOS state: mGPDH-driven respiration is decreased at both temperatures, and CI-driven respiration is increased at 12 °C. The addition of PG did not differentially affect ET-capacity, except when respiration was driven by mGPDH at 12°C in *3XtubAOX* larvae (Supplemental Figure S2C). As expected, AA completely abolished respiration in control larvae, and AOX was still able to sustain 20-28% of ET-capacity respiration in *3XtubAOX* larvae (Supplemental Figure S2D). These results indicate that AOX can contribute to electron transfer, especially via mGPDH, even at low ΔΨm conditions.

During the Leak respiratory state, an intrinsically uncoupled condition achieved by inhibition of the ATP synthase with oligomycin, AOX induced an ∼twofold increase in OCR via CI + mGPDH or via CI only at both temperatures (Figure 3C). Moreover, respiration returned to control levels upon AOX inhibition in *3XtubAOX* larvae, indicating this enzyme is solely responsible for the increased uncoupling of CI-driven OCR (Supplemental Figure S2E). Interestingly, AOX did not alter the Leak respiration driven by mGPDH (Figure 3C), although its inhibition with PG decreased OCR by ∼35% at both temperatures (Supplemental Figure S2E), indicating that uncoupling via mGPDH might be regulated to a particular maximal level. We also examined OCR in the Leak state upon CIII inhibition, and once again, observed complete abolishment of respiration in controls (Supplemental Figure S2F). In *3XtubAOX* larvae, however, the alternative enzyme sustained CI-driven Leak respiration at much higher rates than when CIII was active, an effect more prominently observed at 25 °C (Supplemental Figure S2F). This result is consistent with our previous findings (Othonicar et al., 2024), indicating that AOX only has access to all CoQ reduced by CI when CIII/IV are inactive. Conversely, CIII inhibition caused a drop in mGPDH-driven Leak respiration, which was sustained by AOX at 40-50% of total OCR at both temperatures (Supplemental Figure S2F). This result suggests that CIII/IV and AOX readily share the CoQ molecules reduced by mGPDH.

The data presented above allowed us to calculate the coupling-control ratio (Leak/OXPHOS, (Gnaiger, 2020)) of *Drosophila* larval mitochondria in the presence of AOX. Figure 3D shows that AOX increases OXPHOS inefficiency in both electron transfer pathways tested, regardless of temperature. Notably, mGPDH-driven respiration, which is naturally less coupled than that initiated by CI (Masson et al., 2017), is more affected by AOX, with OXPHOS inefficiency values being twice as high at both temperatures (Figure 3D). This results in greater energy dissipation during mGPDH-driven respiration.

We also analyzed the E-P control efficiency ratio (Gnaiger, 2020) to assess whether the ADP phosphorylation system in mitochondria is exerting any control over the CI-or mGPDH-driven electron transfer pathways to carry out OXPHOS. We observed that at 25 °C, that ratio was low and that it was not altered by AOX, except when respiration was driven by mGPDH, for which the presence of AOX elevated the limitation imposed by the phosphorylation system onto this electron transfer pathway (Figure 3E). At 12 °C, we observed that the OXPHOS limitation was at least twofold higher than at 25 °C for all pathways, but the presence of AOX prevented this limitation from being as high for CI-driven respiration. Interestingly, the E-P control efficiency ratio for mGPDH-driven respiration did not differ from that in the controls at 12 °C, and remained similar to the value at 25 °C (Figure 3E). Taken together, these values indicate that the electron flow initiated by mGPDH in the presence of AOX is either unchanged or more limited by the phosphorylation system, whereas AOX induces a flow that is less limited by the phosphorylation system via CI at 12 °C.

Oxygen consumption increased via CI and decreased via mGPDH, in addition to changes in OXPHOS efficiency and OXPHOS limitation, points to a significant reconfiguration of the larval mitochondrial respiratory system induced by AOX. We then investigated if these changes could be explained by changes in abundance and/or activity of the individual fly OXPHOS components. We first checked transcript levels of *Gpo1*, the coding gene for the ubiquitous form of mGPDH, and observed that at 25 °C no significant differences were observed between control and *3XtubAOX* larvae (Supplemental Figure S1A), as shown in our previous report (Othonicar et al., 2024). At 12 °C, *Gpo1* levels were fourfold higher in the controls, and were kept as low as at 25 °C in *3XtubAOX* larvae. We next examined other components of the larval IMM by blue native-polyacrylamide gel electrophoresis (BN-PAGE) and in-gel activity assays, and detected that the cold suppresses the levels and activity of all forms of respiratory complexes and supercomplexes we identified in control larvae (Figure 3F and G), except those of CII (Supplemental Figure S1D). AOX expression at 12 °C mitigates the cold-induced suppression of the ETS components observed in the control line. The increase in abundance/activity of CI and CI-containing supercomplexes in *3XtubAOX* larvae at 12 °C is consistent with our respirometry data showing increase in CI-driven OCR (Figure 3A-C). In addition, increase in ATP synthase abundance/activity (Figure 3F) appears to support the lower OXPHOS limitation that occurs in *3XtubAOX* larvae at 12 °C (Figure 3E). This agrees with the idea that AOX expression leads to increased electron flow through CI, supported by an increase in ATP synthase activity at cold temperatures.

### Treatments with OXPHOS poisons suggest a key role for CI in the development of AOX-expressing larvae

We next tested if addition of the uncoupler CCCP to the diet could phenocopy the effects of AOX on *Drosophila* development, since AOX can significantly uncouple mitochondria through functional interactions with mGPDH. At 25 °C, increasing concentrations of the uncoupler led to increased body mass in control larvae up to the 20 µM concentration, after which body mass dropped significantly (Figure 4A). The body mass of control larvae with 20 µM CCCP was similar to that of *3XtubAOX* larvae without the uncoupler. AOX larvae were quite sensitive to dietary CCCP, since all concentrations used decreased body masses significantly to levels equivalent to those of controls without uncoupler (Figure 4A). At 12 °C, 20 µM CCCP had no impact on the body masses of control larvae, but was again sufficient to negatively affect those of AOX larvae (Supplemental Figure S1E). In addition, dietary CCCP supplementation induced no significant changes in egg-to-pupa developmental times at 12 or 25 °C (Supplemental Figure S1F, and data not shown). Thus, the previously reported acceleration of ∼9 days in development at 12 °C caused by AOX (Saari et al., 2019a) does not appear to be part of its uncoupling effects.

**Figure 4.**
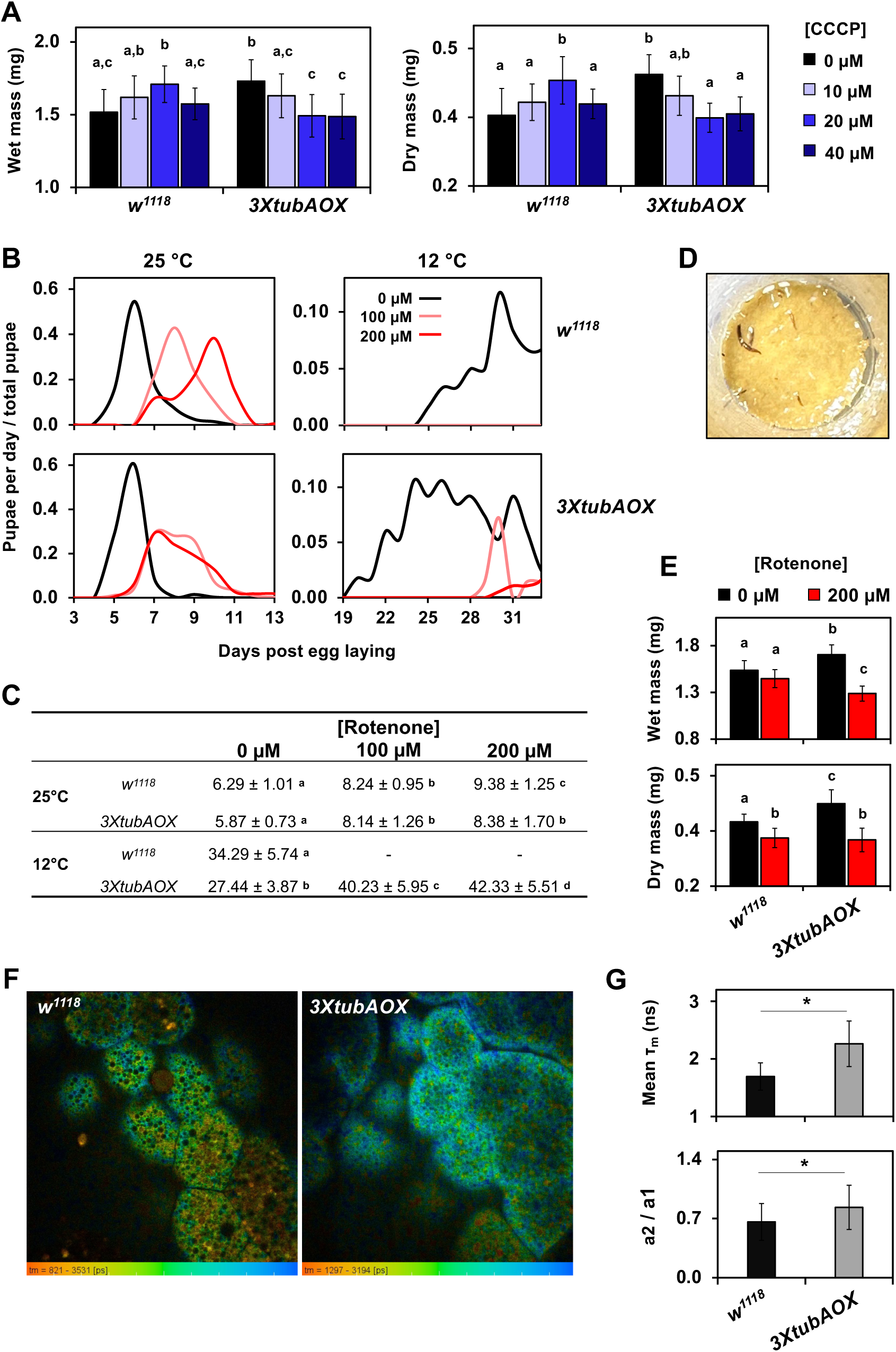
Mitochondrial uncoupling and complex I activity contributes to development of AOX-expressing flies. Wet and dry masses of larvae of the indicated genotype cultured at 25 °C, with dietary supplementation of CCCP (A) or rotenone (E). Data represent means ± standard deviation (SD) from at least 8 replicates, each with 10 larvae. (B,C) Egg-to-pupa developmental time of control (*w^1118^*) and AOX-expressing (*3XtubAOX*) individuals at the indicated rotenone concentration and temperature. Letters ‘a-c’ indicate statistically distinct groups (p < 0.05), according to a two-way ANOVA with Tukey’s *post hoc* test. (D) Representative image showing lethality in *w^1118^* larvae (dark individual larvae on top of diet) at 200 µM rotenone. (F) Representative fluorescence lifetime microscopy images of NADH obtained from fat body cells of wandering L3 larve at 25 °C, with the false-colored scales representing τ, ranging from shorter (red, free NADH) to longer lifetime (blue, enzyme-bound NADH). (G) Mean τ_m_ of free NADH autofluorescence of 5 different samples of larval fat bodies of each genotype. (H) Reverse amplitude rate (a2/a1) of enzyme-bound/free NADH, which provides estimates of the cellular NAD+/NADH ratio.

Given that AOX expression increased larval CI activity and CI-driven mitochondrial oxygen consumption, we then investigated the roles of this complex in development by dietary supplementation with its inhibitor rotenone. Titrating the CI inhibitor at 25 °C revealed a dose-dependent delay in egg-to-pupa developmental time (Supplemental Figure S1G), highlighting a central role for CI in larval growth. In the control line, we observed a delay of 2 and 3 days in development with 100 and 200 µM doses, respectively, relative to the diet without inhibitor. This delay was significantly more modest for the *3XtubAOX* line with 200 µM (Figures 4B and C), suggesting a decreased sensitivity of development to high levels of rotenone. At 12 °C, no control individuals reached the pupa stage with either 100 or 200 µM rotenone, dying at various larval stages (Figures 4B and 4D). In contrast, for the *3XtubAOX* line, the same concentrations of dietary rotenone only delayed pupation by 12–14 days compared to a rotenone-free diet (Figure 4B). Interestingly, the body masses of AOX larvae at 25 °C were more sensitive to dietary rotenone than those of control larvae (Figure 4E). These findings highlight that the increased CI activity in *3XtubAOX* larvae is key in growth and development.

### Higher NAD+/NADH ratio in AOX-expressing larvae is consistent with the metabolic profile of increased growth

The increased CI activity in AOX-expressing larvae may regulate growth metabolism by increasing the rates of mitochondrial NADH reoxidation, therefore increasing the cellular NAD+/NADH ratio. This would drive forward the glyceraldehyde-3-phosphate dehydrogenase reaction of glycolysis, and the NAD+-dependent reactions of the tricarboxylic acid (TCA) cycle and other pathways. To test this, we used fluorescence-lifetime imaging microscopy to quantify the autofluorescence of NADH *in situ* in larval tissues (Figure 4F). The increased lifetime of enzyme-bound NADH autofluorescence and the ratio between enzyme-bound and free NADH (a2/a1) in *3XtubAOX* at 25 °C (Figure 4G and H, Supplemental Figure S3A) are consistent with our hypothesis that AOX induces elevated NAD+/NADH ratio via increased CI activity.

We next sought to evaluate the impacts of AOX expression on the larval metabolome. Using targeted metabolomics based on gas chromatography-mass spectrometry (GC-MS) to quantify key metabolites in wandering L3 larvae at 25 °C, we identified by Principal Component Analysis (PCA) a clear separation between samples from the control and *3XtubAOX* lines, indicating significant alteration of the metabolic profile caused by AOX (Figure 5A). This was primarily due to a substantial decrease in the levels of most metabolites (Figure 5B), such as intermediates of the TCA cycle, notably citrate, fumarate and malate (Figure 5C). Among the amino acids, while glutamate and methionine were more abundant in the presence of AOX, the levels of most other amino acids were decreased, with aspartate showing a notable drop of ∼60% (Figure 5C). Levels of G3P, the substrate of mGPDH, also decreased in *3XtubAOX* larvae (Figure 5B). We also investigated the larval metabolome at the mid-L2 stage, during which rapid growth occurs. Additionally, we included in the analysis two fly lines expressing lower levels of AOX, *tubAOX^35^* and *tubAOX^7^* (Kemppainen et al., 2014; Saari et al., 2017) and focused on key TCA cycle metabolites, G3P, lactate, and 2-hydroxyglutarate (2-HG; note that the larval 2-HG pool is primarily composed of L-2-HG (Li et al., 2017)). Once again, PCA revealed a clear separation between the *3XtubAOX* and *w^1118^* samples, with samples from the *tubAOX^35^*and *tubAOX^7^* lines showing intermediate metabolic profiles (Figure 5D and E). This suggests a dose-dependent effect of AOX, with higher AOX expression leading to a more pronounced decrease in the abundance of the selected metabolites. The separation between the larval metabolic profiles of *w^1118^*and *3XtubAOX* was also evident in broader, untargeted metabolomics analyses based on liquid chromatography-mass spectrometry (LC-MS) for L2 and L3 larvae, again with intermediate profiles for *tubAOX^35^* and *tubAOX^7^*, as analyzed by Partial Least Squares Discriminant Analysis, PLS-DA (Supplemental Figure S3B).

**Figure 5.**
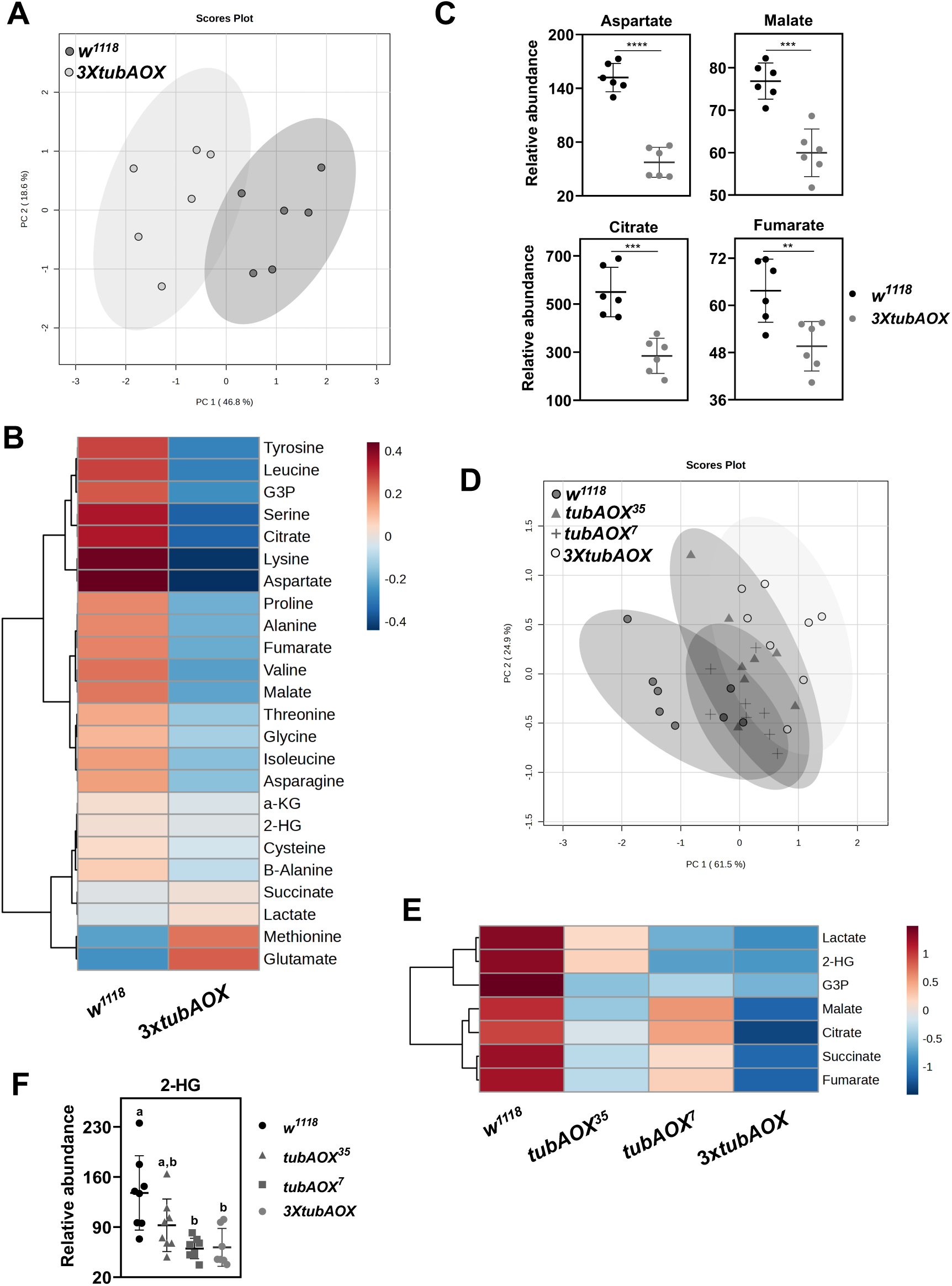
Targeted metabolomics indicates AOX-induced difference in the larval metabolism. Levels of the indicated metabolites were determined by gas chromatography-mass spectrometry (GC-MS) to assess relative abundances among control (*w^1118^*) and AOX-expressing lines. The data was examined by principal component analyses (PCA) to show the overall differences between *w^1118^* and *3XtubAOX* L3 larvae (A), and among *w^1118^*, *tubAOX^35^*, *tubAOX^7^* and *3XtubAOX* mid-L2 larvae (D). The heat maps (B, E) and scatter plots (C, F) depict relative levels of the indicated metabolites. **, *** and **** indicate, respectively, significant differences with p < 0.01, p < 0.001 and p < 0.0001) determined by Student’s t-tests, and letters ‘a-c’ denote differences according to one-way ANOVA with Tukey’s *post hoc* test (p < 0.05). 2-HG, L-2-hydroxyglutarate; α-KG, α-ketoglutarate; G3P, glycerol-3-phosphate.

Although not statistically significant, the LC-MS data for L3 larvae showed a tendency of increased NAD^+^ levels and NAD+/NADH ratios in the AOX-expressing lines in a dose-dependent manner (Supplemental Figure S3C), which is in agreement with our *in situ* NADH quantification data (Figure 4F-H). We also observed a tendency in AOX-expressing lines of elevated ATP levels and ATP/ADP ratios (Supplemental Figure S3D), which is consistent with the increased ATP synthase activity shown in Figure 3. We then explored all glycolytic and TCA cycle metabolites identified in the LC-MS data to see the impacts of higher NAD+ and ATP levels on central carbon metabolism of the larvae. Compared to *w^1118^* controls, the levels of almost all of these metabolites in *3XtubAOX* larvae were decreased, whereas phosphoglycerate (2/3-PG), phosphoenolpyruvate (PEP) and α-ketoglutarate (α-KG) were elevated (Supplemental Figure S3E). This suggests that AOX generally increases the flux of the larval proliferative metabolism, in agreement with our biomass accumulation data of Figure 2, with the buildup of the key metabolites PEP and α-KG. In addition, the lower levels of lactate in AOX-expressing larvae are consistent with enhanced mitochondrial oxidative metabolism (Supplemental Figure S3E). We also observed lower levels of 2-HG, a result consistent with the fact that the majority of this pool consists of L-2-HG, whose production by lactate dehydrogenase (Ldh) is dependent on NADH (Figure 5F). Despite the lower levels of lactate and L-2-HG, *Ldh* transcript levels at 25 °C are elevated in *3XtubAOX* (Supplemental Figure S1A). Altogether, our metabolomics data supports a model in which AOX expression accelerates larval growth by increasing oxidative metabolic flux, which in turn is caused by the elevated CI-driven NAD+/NADH ratios.

## Discussion

Under stressful cold temperatures, the xenotopic expression of *C. intestinalis* AOX in *Drosophila* significantly accelerates development and enhances larval viability (Saari et al., 2019a). In this study, we set out to investigate how AOX expression leads to this phenotype, considering the substantial impact that thermal variation has on insect developmental metabolism, physiology and behavior (Chen et al., 2014; Ghosh et al., 2013; McParland et al., 2015). Using infrared thermographic imaging and chip calorimetry, we showed that whole-body temperature and heat production in AOX-expressing larvae and pupae are not significantly different when compared to the *w^1118^* controls. However, the signal in the real-time readings of the calorimetry experiments indicates that the dynamics of body heat production in AOX and control animals are not the same. In the larvae, AOX induced an increase in the signal noise, which was associated with increased larval mobility, whereas in the immobile pupae, ∼10 s-long heat bursts occurred in ∼1 min cycles. Interestingly, these heat bursts are also observed for the control pupae, although much more infrequently. To our knowledge, this phenomenon has not yet been characterized and obviously needs to be further investigated. At the level of the organelle, the expression of AOX in mammalian and *Drosophila* cells in culture does increase local mitochondrial temperature, followed by a cooling period that maintains cellular thermal homeostasis (Jacobs et al., 2024; Terzioglu et al., 2023). Perhaps this cooling can also maintain organismal thermal homeostasis, at least in *Drosophila*.

Our mitochondrial physiology studies suggest that AOX significantly increase the inefficiency of larval OXPHOS, primarily via mGPDH-linked respiration. This appears to be due to a functional interaction between mGPDH and AOX, which is futile for ATP production, as neither enzyme contributes to ΔΨm. This larval ETS configuration constitutes a molecular arrangement similar to the thermogenic mechanism observed in plants like the skunk cabbage *Symplocarpus renifolius* (Ito et al., 2004; Onda et al., 2008) and the lotus flower *Nelumbo nucifera* (Watling et al., 2006), in which a non-proton-pumping dehydrogenase and a terminal oxidase enable mitochondrial uncoupling and thermogenesis. The mGPDH-AOX pathway in AOX larvae is functionally important at 12 °C, because it sustains mGPDH-driven Leak respiration at levels comparable to control larvae, despite the decrease in mGPDH-driven OXPHOS respiration that is induced by AOX expression. Furthermore, at this temperature, AOX actively supports approximately one third of the OXPHOS respiration via mGPDH, as judged by the combination of treatments with the AOX inhibitor PG and the CIII/IV inhibitor AA. Interestingly, it has been suggested that mGPDH is less cold-sensitive than other IMM dehydrogenases, being able to support preheating of the flight muscles in the bumblebee *Bombus terrestris* when ambient temperatures approach freezing (Masson et al., 2017).

With increased mitochondrial uncoupling and OXPHOS inefficiency, we predicted that AOX expression could compromise larval body mass accumulation, despite accelerating development, since decreased energy conservation for ATP synthesis would oppose the intense larval anabolic metabolism. However, we observed a significant increase in biomass at both temperatures, a feature that was more pronounced under cold thermal stress. This in fact explains the elevated heat capacity and exchange rates of AOX larvae. We noticed AOX caused upregulation of larval food consumption, a behavior that allowed the larvae not only to counteract the excess energy dissipation, but also to store extra biomass energy as fat. This was unexpected, but it might indicate a mechanism that could sense lower rates of energy conversion for larval biomass storage due to mild mitochondrial uncoupling, and prepare the organism for this elevated uncoupling during the pupal stage, when it relies exclusively on stored energy for the intense metabolic processes of metamorphosis. This is consistent with the results of our developmental assays in the presence of the mitochondrial uncoupler CCCP, which showed increase in body mass of control *w^1118^* larvae up to 20 μM of dietary CCCP. This elevated larval storage capacity appears to be dependent on a nutrient-rich diet, since we have shown previously that a different AOX-expressing fly model has high rates of pupal lethality when the larvae are cultured on a low-nutrient diet (Saari et al., 2019b, 2019a). The body weight of *3XtubAOX* larvae were also sensitive to the uncoupler, decreasing as the concentrations of dietary CCCP increased at 25 °C. In addition to the fact that control larvae also showed decrease in body weight with 40 μM of dietary CCCP, our data indicates that AOX expression may have a hormetic effect on larval mass accumulation and that the levels of AOX in the *3XtubAOX* line are about optimal for this feature. In fact, AOX expression based on the *UAS*/*GAL4* system, which leads to much higher levels of AOX, does cause lower larval weight and fat accumulation (Camargo et al., 2021).

Nevertheless, mitochondrial uncoupling alone does not seem to explain all developmental phenotypes caused by AOX expression in the *3XtubAOX* line. Although 20 μM of dietary CCCP at 12 °C affects biomass accumulation of AOX-expressing larvae, it does not accelerate development of this line, nor does it alter any parameters of the control animals. We also observed an increase in CI-driven OXPHOS respiration in AOX larvae, which was associated with an increase in ATP synthase activity at 12 °C. This indicates higher electron flux via CI and consequently enhanced NADH reoxidation. We in fact showed that *3XtubAOX* larvae have a higher cellular NAD+/NADH ratio, which was associated with lower levels of most TCA cycle intermediates. This supports the idea that these intermediates are being rapidly consumed, i.e., that the metabolic flux through the TCA cycle is elevated, stimulated by its NAD+-dependent reactions, and requiring greater oxidation of acetyl-CoA and/or other metabolites. Whereas the development of AOX-expressing larvae is more resistant to the CI inhibitor rotenone, especially at 12 °C, biomass accumulation appears to be more sensitive to such poison, highlighting the importance of CI for the larval proliferative metabolism driven by AOX. The increase in CI-driven respiration is at least in part due to elevated levels of CI-containing supercomplexes, which may be streamlining electron transfer from NADH to oxygen to optimize OXPHOS in an attempt to counterbalance the AOX-induced uncoupling. We have shown previously that levels of supercomplexes correlate with mitochondrial DNA copy number and development in diverse lines originated from natural *D. melanogaster* populations (Salminen et al., 2017). In addition, our data of CI-driven Leak respiration shows that CIII inhibition with AA stimulates OCR via AOX to levels more than twofold higher, reaching values comparable to the levels of AA-resistant respiration in the OXPHOS state (when ATP synthase is not inhibited), consistent with our recently published data (Othonicar et al., 2024). This indicates AOX does not oxidize most CI-reduced CoQ molecules in larval mitochondria, perhaps because these are trapped inside supercomplexes and only become available to AOX upon CIII inhibition. It is interesting how AOX appears to be excluded from electron transfer within larval CI-containing supercomplexes and yet makes the larval biomass more sensitive to CI inhibition; this certainly warrants further investigation.

As main regulators of cellular and systemic NAD+/NADH ratio, the mitochondrial status largely influences cell and organismal metabolism. It is yet to be fully understood how mitochondria match their biosynthetic functions with the metabolic requirements of tissue growth (Suomalainen and Nunnari, 2024), but our data points to important roles for both mild uncoupling and NADH reoxidation. Since the cataplerotic reactions of the TCA cycle are critical for larval growth (Jacobs et al., 2020; Owen et al., 2002), we hypothesize that an increase in TCA cycle activity in AOX-expressing larvae favors the utilization of its intermediates for biosynthetic pathways, with increased oxidative mitochondrial metabolism. This effect is similar to the metabolic adaptations seen in *D. melanogaster* larvae mutant for the *Dg* gene (the ortholog of the mammalian *dystroglycan 1* gene), which exhibit greater cold tolerance also due to increased mitochondrial metabolism (Takeuchi et al., 2009). Often, the larval proliferative metabolic program is compared to that of cancer cells, characterized by the utilization of glycolysis to produce large amounts of ATP, even under aerobic conditions, a phenomenon called aerobic glycolysis or Warburg effect (Drummond-Barbosa and Tennessen, 2020; Vander Heiden et al., 2009; Warburg, 1956). Even so, mitochondrial respiration is fundamental for maintaining the TCA cycle and cataplerosis. The relevance of OXPHOS becomes more evident when the inhibition of respiratory complexes leads to the cessation of tumor cell proliferation (Bianchi et al., 2018; Galber et al., 2020; Tan et al., 2019; Wang et al., 2021). However, it has been shown in osteosarcomas that the decrease in proliferation caused by ATP synthase inhibition is reversed by the addition of mitochondrial uncouplers (Galber et al., 2020; Sullivan et al., 2015). This suggests that the role of OXPHOS in ATP production is not the most relevant for cell proliferation, but rather it is the electron flow through the ETS and, consequently, the reoxidation of NADH or other reduced metabolites (Galber et al., 2020). Most ATP is then produced via aerobic glycolysis.

Our data does not exactly sustain the idea that the growth acceleration caused by AOX expression is due to increased aerobic glycolysis, at least not in the traditional view of the phenomenon. Although we observed lower levels of most glycolytic intermediates in AOX larvae, which in combination with higher NAD+/NADH ratio could indicate increased metabolic flux through this pathway, the expected increase in lactate levels by aerobic glycolysis was not observed. On the contrary, lactate levels were lower despite higher *Ldh* transcript levels in the presence of AOX at 25 °C; at 12 °C, *Ldh* levels were as low as in the controls. In addition, lower levels of pyruvate and an accumulation of PEP and 2/3-PG suggest that the ATP-producing pyruvate kinase reaction of glycolysis is downregulated. The increased activity of the mitochondrial ATP synthase in AOX-expressing larvae must then be fulfilling the ATP requirements of the larval proliferative metabolism. Lower levels of pyruvate, in combination with lower oxaloacetate, citrate, succinate, fumarate and malate, are consistent with increased flux through the TCA cycle in the presence of AOX. The accumulation of α-KG, however, is interesting, as it may indicate downregulation of the TCA cycle enzyme α-KG dehydrogenase and/or upregulation of glutamate dehydrogenase (GDH) or glutamate-oxaloacetate transaminase 2 (GOT2). *D. melanogaster* larvae also naturally produce high concentrations of L-2-HG by oxidation of NADH and reduction of α-KG, catalyzed by Ldh or the TCA cycle enzyme malate dehydrogenase (MDH2) (Du and Hu, 2021; Li et al., 2017; Mahmoudzadeh et al., 2020). We demonstrated that although AOX larvae grow more, L-2-HG abundance is significantly decreased in the L2 stage, which is also consistent with the increase in NAD+/NADH ratio and with the elevated levels of α-KG. In fact, hypoxia and other forms of mitochondrial ETS impairment, with decreased NADH reoxidation, correlate with the accumulation of larval L-2-HG (Li et al., 2018; Mahmoudzadeh et al., 2020).

*De novo* fatty acid anabolism is dependent on transport of citrate from the mitochondria to the cytosol, where the enzyme citrate lyase cleaves it into oxaloacetate and acetyl-CoA, the latter needed for the rate-limiting step in fatty acid synthesis, the acetyl-CoA carboxylase reaction (Icard et al., 2021). The knockdown of citrate lyase in flies causes lethality at the larval stage (Saari et al., 2019b). In AOX larvae, the low levels of the lipid precursor citrate is in agreement with the higher levels of lipids stored. Interestingly, we speculate that, in addition to the reaction catalyzed by the TCA cycle enzyme citrate synthase, a substantial portion of the citrate in AOX larvae is being synthetized by carboxylation of α-KG via activity of another TCA cycle enzyme, isocitrate dehydrogenase (IDH), forced by the accumulation of this metabolite. Citrate plays a central role in the proliferation of cancer cells, due to the rapid turnover of oxaloacetate and acetyl-CoA, and α-KG carboxylation from glutamine metabolism, ensuring high cataplerosis for tumor growth (DeBerardinis et al., 2008; Icard et al., 2021; Jiang et al., 2023; Wise et al., 2011). It would be interesting to test if combining AOX expression with modulation of IDH levels or expression of mutant forms of the enzyme in the *D. melanogaster* larvae could have a further impact on biomass accumulation and growth. Because we showed here the impact of AOX expression on the electron transport pathway initiated by mGPDH, we can speculate that electron transport initiated by other FAD-linked IMM dehydrogenases, which do not form supercomplexes, is also impacted similarly. A decrease in CII activity analogously to that seen for mGPDH could also explain the elevated levels of α-KG, although our in-gel activity assays show no alterations in CII in AOX-expressing larvae. The inability of fly mitochondria to import succinate into the matrix (Tulp et al., 1971; Van Den Bergh and Slater, 1962) and data not shown) certainly prevents us from testing CII-linked OCR via classical mitochondrial physiology studies.

The increase in cellular NAD+/NADH ratio by mitochondrial CI in AOX-expressing larvae implies that the mitochondrial NADH shuttle systems are operating at higher rates. The export of citrate from mitochondria also serves the purpose of importing malate via the malate-citrate antiporter of the IMM. Cytosolic malate may be formed by reduction of oxaloacetate generated by citrate lyase, with consumption of cytosolic NADH. Once in the mitochondrial matrix, malate is converted back into oxaloacetate by MDH2 with production of NADH, the substrate of CI (Guo et al., 2023; Icard et al., 2021). The malate-aspartate shuttle combines the mitochondrial entry of malate with the export of α-KG, at the same time that another antiporter imports glutamate and exports aspartate. In the cytosol, aspartate and α-KG are used in a transamination reaction to form oxaloacetate and glutamate. The latter is reduced by cytosolic NADH to malate, which upon entry to mitochondria is oxidized back to oxaloacetate to generate intramitochondrial NADH. The cycle is completed by transamination of glutamate and oxaloacetate to form aspartate and α-KG by the mitochondrial enzyme GOT2. Low levels of aspartate and high levels of glutamate and α-KG, as seen in the AOX-expressing larvae, are indicative of altered functioning of this shuttle (Borst, 2020). The malate-citrate and the malate-aspartate shuttles become even more important for cellular NADH reoxidation if we consider that the activity of a third shuttle, the G3P shuttle, is decreased upon larval AOX expression, as observed by the lower rates of G3P oxidation linked to oxygen consumption, which is initiated by mGPDH. G3P is formed by the cytosolic GPDH via reduction of the glycolytic metabolite dihydroxyacetone phosphate (DHAP) with consumption of NADH. In the intermembrane side of the IMM, mGPDH converts G3P back into DHAP and transfers the electrons directly to CoQ, and then to CIII/IV (Mráček et al., 2013). Our metabolomics show that larval G3P levels are low in AOX larvae, which is consistent with the increase in its relative fat mass, since G3P is required for the triacylglycerol esterification reactions. Higher cGPDH activity and/or lower mGPDH activity are associated with increased lipid synthesis and triacylglycerol levels in other organisms (Dulermo and Nicaud, 2011). It is yet to be shown for AOX larvae if the decrease in mGPDH-driven OCR stimulates lipid synthesis by not competing for G3P consumption, or if the high utilization of G3P in lipid synthesis makes this metabolite limiting, regulating mGPDH activity.

In summary, by expressing AOX during the growth phase of *Drosophila*, we were able to observe a stimulation of mitochondrial metabolism, with a clear contribution to the larval anabolic processes, which were especially important at stressfully cold temperatures. The larval proliferative metabolism benefited from the mild uncoupling generated by the mGPDH-AOX functional interactions and the increased rate of NADH reoxidation by CI and CI-containing supercomplexes. It is possible that the changes in the ETS configuration we have shown here are accompanied by changes in mitochondrial morphology that support the increased metabolism, but this is also yet to be studied. In addition, the higher rates of mitochondrial NADH reoxidation suggest that larval growth is at least in part naturally limited by the capacity of their mitochondria to import reduced NADH equivalents from the cytosol for reoxidation, providing in exchange ATP and precursors for macromolecule biosynthesis. In that sense, it is not surprising that Ldh and the elevated production of lactate are so important for larval growth (Li et al., 2017), as this contributes to cellular NADH reoxidation. However, in proliferating human cells in culture, aerobic glycolysis and lactate production do not correlate with growth and appear to occur because of the saturation of the mitochondrial NADH shuttles (Wang et al., 2022). Whether the cytosolic Ldh reaction remains important upon the stimulation of mitochondrial metabolism caused by larval AOX expression, and whether AOX expression causes similar metabolic changes in other proliferative systems, such as tumor cells, are also interesting avenues of research to pursue.

## Materials and Methods

### Fly lines and culture

The fly lines *w^1118^* (genetic background), *tubAOX^35^* (single copy of *C. intestinalis* AOX transgene under the control of the *αTub84B* gene promoter, inserted into chromosome X), *tubAOX^7^* (chromosome 3), and *3XtubAOX* (three copies of AOX transgene, in chromosomes X, 2 and 3) (Kemppainen et al., 2014) were originally provided by Dr. Howard T. Jacobs’s laboratory (Tampere University, Finland). The line *UAS-empty^2nd^*; *UAS-empty^3rd^* was created via standard crosses using balancer chromosomes and the original lines *UAS-empty^2nd^*(empty pUASTattB vector, inserted in chromosome 2) and *UAS-empty^3rd^*(chromosome 3) (Andjelković et al., 2015). The flies were cultured under controlled temperatures (25 °C or 12 °C) and a 12-hour light/12-hour dark photoperiod on a standard diet, as described elsewhere (Fernandez-Ayala et al., 2009). For data in Figure 4 and Supplementary Figure S1, the diet was supplemented with the mitochondrial uncoupler CCCP or the CI inhibitor rotenone by adding 40 µl of an ethanol solution to reach the final concentration indicated in the figure legends. Vials containing approximately 6 ml of diet, 20 female flies, and 10 male flies were maintained at 25 °C for mating and egg laying for ≤ 24 hours, after which, adult flies were removed, the embryos allowed to develop at 25 °C or 12 °C until reaching the indicated stage, and the individuals randomly collected for experimentations. Alternatively, to assess the egg-to-pupae developmental time, vials were monitored daily to track the appearance of pupae, and the number of pupae per vial was recorded on successive days relative to the day of egg laying.

### RNA extraction and quantitative PCR analyses

Four wandering L3 larvae (∼120 h post-oviposition) were collected, rinsed, fast-frozen in liquid nitrogen, and homogenized in 300 μl of TRI Reagent™ Solution (Invitrogen™). Following the addition of 120 μl chloroform, samples were vortexed for 15 s, incubated at room temperature for 15 min, and centrifuged at 12,000 x*g* for 20 min. The supernatant was transferred to a new tube, mixed with 300 μl ice-cold isopropanol, and centrifuged again. RNA pellets were washed twice with 500 μl ice-cold 75% ethanol, centrifuged at 12,000 ×g for 10 min, resuspended in 30 μl RNase-free water, and stored at-80 °C. cDNA synthesis was performed using the High-Capacity cDNA Reverse Transcription Kit (Applied Biosystems™) with 500 ng of total RNA, following the manufacturer’s instructions. Quantitative real-time PCR was conducted on the CFX Opus Real-Time PCR System (Bio-Rad) using PowerUp™ SYBR™ Green Master Mix (Applied Biosystems™). Relative transcript levels were determined using the ΔΔCt method with the following primers: for *Gpo1* 5’TCCTCGGTGGGCGTTGGATT3’ and 5’CGACGGCCGCGTCATCTTCT3’; for *Ldh* 5’ATACACCTCCTGGGCCATTG3’ and 5’CAATGCCATGTTCGCCCAAA3’; for *AOX* 5’ATTTTCTTTGGCTTACTTAATCTCAC3’ and 5’CAATTTCTGGCGCTTTCA3’; and for the housekeeping *eIF-1A* 5’ATCAGCTCCGAGGATGACGC3’ and 5’GCCGAGACAGACGTTCCAGA3’.

### Larval mobility, size, biomass and food intake assessments

Larval mobility assays were adapted (Nichols et al., 2012), and details described in Garcia et al. (Garcia et al., 2021). Briefly, the larvae were collected, rinsed in deionized water, placed on 2% agar plates with a millimetric grid scale (0.2 cm²), and allowed to acclimate for 5 minutes, after which speed of locomotion was quantified by counting the number of lines crossed per minute. The number of complete peristaltic contractions (from anterior to posterior end of the larva) per minute was recorded manually under a stereoscopic microscope. In addition, the standard deviation of the signal noise of the calorimetry real-time readings (σ) was used as another proxy of mobility, as the larvae were able to move inside the calorimetry channels.

To quantify body mass, groups of 10 individuals (24-30 technical replicates, 3 biological replicates, ∼800 individuals from each line) were randomly collected, rinsed in deionized water, dried on Kimwipe tissue, and placed on pre-weighed 1.5 ml microcentrifuge tubes on a precision balance with an accuracy of 0.0001 g (Shimadzu, ATX224). Samples were then dried in a dry bath at 65°C until the weight stabilized (at least 2 hours) and tubes weighted again to obtain the dry mass. Water content was calculated subtracting the dry mass from the wet mass. The samples were then incubated with 1 ml of ethyl ether for 24 hours, discarded and the process repeated 3 times, after which the samples were dried at 65°C again, as described earlier. The dried, fat-free larvae were transferred to new tubes and weighted again to determine lean mass and fat mass (dry mass - lean mass). The values per tube were divided by 10, generating an average per individual larva. Total protein content were estimated using the Bradford method (Bradford, 1976) with homogenates of sets of 20 larvae (in 14-15 independent experiments) macerated on ice with 1 ml of isolation buffer (250 mM sucrose, 5 mM Tris-HCl pH 7.5, 2 mM EGTA, pH 7.4) and a glass homogenizer. Individual larval protein content was normalized by the lean mass and the value per tube divided by 20.

For larval body dimensions, individuals were collected and processed as described earlier, and carefully immersed in ∼80°C water for ∼3 s to achieve complete immobilization and body distension. They were then photographed under a microscope with an attached camera, and the images were analyzed using the ImageJ software (http://imagej.nih.gov/ij/), where size was estimated by measuring the length and width of the larval body, represented by longitudinal and transversal lines. At least 72 larvae from each line were sampled at each tested temperature, with 3 biological replicates.

Food consumption assays were adapted from (Kaun et al., 2007). Briefly, larvae were cultured post-oviposition for 84-90 hours at 25 °C, or for 20-30 days at 12°C, collected and processed as described earlier, and placed on Petri dishes with Kimwipe tissues soaked with 1% PBS for a 3 hour fasting period. They were then transferred to plates containing a yeast paste (32.5% w/v in water) dyed with Brilliant Blue R (0.7% w/w) and allowed to feed for 5 minutes at 25°C or for 30 minutes at 12°C, after which they were immediately immobilized in ∼80°C water for 5 s, and photographed in the ventral position using a microscope with an attached camera. The images were analyzed using Image J to calculate the blue area (representing food consumption) per total larval area.

### Infrared thermography and calorimetric measurements

A chip flow calorimeter was modified (Lerchner et al., 2024) based on previous versions (Hartmann et al., 2014; Wolf et al., 2015) to measure continuous body heat production of wandering L3 larvae and prepupae. Essentially, the equipment uses a silicon chip with thermopiles surrounded by two thermostats, which serve as thermal power detectors. Due to its low thermal time constant (24 s), short-term changes in heat production rate can be detected. Individuals were weighed and positioned in segments between two nodes of a 0.2 mm polyamide fiber containing 1% phosphate-buffered saline (PBS), which allowed sample transport through the 1.8 mm-wide calorimeter flow channel. Each experiment was carried out using two individuals at a time, with measurements obtained sequentially in cycles lasting approximately 18 minutes each. A segment containing an inert spongy material was used as reference.

A thermographic camera sensitive to infrared radiation (Flir SC660, Switzerland) was employed to detect surface body temperature and to infer heat production. For each experiment, larvae or prepupae cultured at 25 °C were collected and transferred to Petri dishes (carefully dried to avoid interferences of evaporation with the temperature measurements) covered with anti-reflective masking tape (Emissivity = 0.95) inside an incubator with controlled temperature of 12 °C. The camera was fixed in a tripod inside the incubator, which was open for only ∼5 s for positioning of the Petri dishes, not allowing significant changes in the temperature inside the incubator.

Simultaneous thermal image capture of animals of both genotypes was performed at every 5 s for ∼8 min, time required for the individual body temperatures to reach 12 °C.

The data was analyzed using Therma Cam Researcher 2.10 software. Nonlinear regressions of body temperature decay as a function of time were calculated using GraphPad Prism, v. 8.0.1. Larval heat capacities were estimated based on the equation: [water content (mg) X 4.18 mJ.K^-1^.mg^-1^] + [fat content (mg) X 2.00 mJ.K^-1^.mg^-1^] + [protein content (mg) X 1.40 mJ.K^-1^.mg^-1^], where 4.18, 2.00 and 1.40 mJ.K^-1^.mg^-1^ represent, respectively, the specific heat capacities of water, fat and protein. Heat exchange rates were calculated as [heat capacity X 0.26 K/s (cooling rate derived from Figure 1D)].

### Confocal and fluorescence-lifetime imaging microscopy

Fat bodies from wandering L3 larvae were dissected in cold 1X PBS and immediately fixed in 4% paraformaldehyde for 40 minutes. Following two washes, the samples were incubated with BODIPY (1:500), DAPI (1:4000), and phalloidin-TRITC (1 μg/ml, Sigma, USA) to stain lipid droplets, nuclei, and actin filaments, respectively. After three additional washes, the samples were mounted in Fluoromount Aqueous Mounting medium (Sigma, USA) and imaged using a Leica SP8 confocal microscope and the LAS X software. Images were analyzed using ImageJ to calculate the lipid droplets area.

Fluorescence-lifetime images were acquired using an inverted confocal microscope (LSM 780-NLO Zeiss Axio Observer Z.1, Carl Zeiss AG, Germany) equipped with a time-correlated single-photon counting (TCSPC) module (Becker & Hickl) and a Chameleon Discovery Nx pulsed laser (100 fs pulse duration, 80 MHz repetition rate; Coherent Inc., Santa Clara, CA), housed at the National Institute of Science and Technology on Photonics Applied to Cell Biology (INFABIC), University of Campinas (Unicamp), São Paulo, Brazil. The larval fat bodies were dissected as described above and the tissues were excited at a wavelength of 760 nm with an average power of 20 mW at the objective’s output. The laser was focused on the sample using an EC Plan-Neofluar 40×/1.3 oil-immersion objective (Carl Zeiss AG, Germany). The autofluorescence signal of NADH was detected by a photon-counting photomultiplier tube (PMT) (Becker & Hickl, SPC-830) after passing through a cube containing a BandPass filter (445/45 nm), forming an image with a field of view of 512 × 512 pixels. The TCSPC electronic system (Becker & Hickl, SPC-830) enabled the direct acquisition of fluorescence intensity images via two-photon excitation fluorescence (TPEF), which were subsequently processed by SPCImage software (Becker & Hickl) to generate time-domain fluorescence-lifetime images. A shortpass 690 filter was used to block the excitation laser at the detector input, allowing only the TPEF signal to pass through.

Images were analyzed using SPCImage to calculate the decay matrix for regions of interest (ROIs) and generate the decay parameters. A multiexponential decay was calculated for two components using the Maximum Likelihood Method, yielding a1 and a2 (%) and t1 and t2, as well as the mean lifetime for each ROI.

### Mitochondrial oxygen consumption experiments

Mitochondrial physiology was assessed through real-time oxygen consumption measurements using the high-resolution respirometry system Oxygraph-2k (O2k, Oroboros Instruments), according to Garcia et al. (Garcia et al., 2021). Total homogenates of L3 larvae cultured at 25 or 12°C were prepared in 1 ml of isolation buffer and assayed at the respective culture temperature. One or two hundred μl of the homogenate were added to the chambers containing respectively 1900 or 1800 μl of assay buffer (120 mM KCl, 5 mM KH_2_PO_4_, 3 mM Hepes, 1 mM EGTA, 1 mM MgCl2, 2% BSA, pH adjusted to 7.4) at stirring of 750 rpm. Oxidizable substrates/inhibitors were added separately to analyze different electron transfer pathways: 5 μl of 2 M pyruvate (final concentration of 5 mM) and 7.5 μl of 0.4 M malate (1.5 mM) for CI; and 80 μl of 0.65 M G3P (26 mM) and 1 μl of 0.01 M rotenone (0.005 mM) for mGPDH.

The OXPHOS respiratory state was estimated after addition of 4 μl of 0.5 M ADP (1 mM); the following titration of 1 μl of 8 mM CCCP (4 μM per dose) allowed for maximal electron transfer capacity (ET-capacity state). The Leak state was achieved in independent experiments with addition of 1 μl 25 μM oligomycin (12.5 nM). For each respiratory state, oxygen consumption exclusively supported by AOX was examined by adding 2 μl of 0.05 M antimycin A (0.05 mM), followed by titration of 2 μl of 0.1 M propyl gallate (0.1-0.2 mM); the opposite order of inhibitors were used independently to obtain CIII/IV-supported oxygen consumption. OCR were calculated by subtracting the residual respiration after inhibition of AOX, CIII and CI, and were normalized by the total protein content of larval homogenates, obtained by the Bradford method.

The relative limitation rate of the ETS by the ADP phosphorylation system, which indicates the capacity of ETS unused for OXPHOS in each experimental condition, was calculated using the formula 1 – (OXPHOS OCR/ET-capacity OCR). Uncoupling of oxygen consumption from ADP phosphorylation (OXPHOS inefficiency) was estimated via the coupling-control ratio (Leak OCR/OXPHOS OCR) (Gnaiger, 2020).

### Blue native-polyacrylamide gel electrophoresis and in-gel activity assays

Crude mitochondrial extracts were prepared from 120 wandering L3 larvae (∼120 h post-oviposition) using protocols adapted from elsewhere (Murari and Owusu-Ansah, 2021; Timón-Gómez et al., 2020). Larvae were homogenized in 700 μl ice-cold preparation buffer (250 mM sucrose, 10 mM Tris HCl, 0.15 mM MgCl_2_, pH 7.4, 1X Halt™ Protease Inhibitor) and centrifuged at 4 °C for 5 min at 600 x*g*, followed by supernatant discard and another centrifugation of 9000 x*g* for 10 min. Pellets were resuspended in 300 μl isolation buffer, and protein quantification was performed according to the Bradford method. Approximately 50 μg of mitochondrial proteins were recentrifuged at 9000 x*g* for 10 min and resuspended in 20 μl 1X NativePAGE™ Sample buffer containing 1.5% digitonin. After incubation on ice for 20 min, samples were centrifuged at 20,000 x*g* for 20 min, and 1.5 μl Coomassie Blue G-250 was added. Samples were then electrophoresed on 3-12% NativePAGE™ gels using NativePAGE™ Anode and Cathode Buffers at 4-8 °C for 30 min at 150 V using light blue cathode buffer, followed by 150 min at 200 V using dye-free cathode buffer.

Gels were stained with Colloidal Blue or used for in-gel activity assays. For identification and quantification of active CI complexes and supercomplexes, 0.1 mg/ml NADH and 2.5 mg/ml nitro blue tetrazolium (NBT) in 2 mM Tris-HCl (pH 7.4) for 1-2 h were used, whereas for CII, 20 mM sodium succinate, 2.5 mg/ml NBT and 0.2 mM phenazine methosulfate in 5 mM Tris-HCl (pH 7.4) for 3-4 h were required. CIV assays were performed with 0.05 mM cytochrome c and 0.05% diaminobenzidine in sodium phosphate buffer (pH 7.2) for 3-4 h, and ATP synthase assays with 14 mM magnesium sulfate, 0.2% lead nitrate and 8 mM ATP in 35 mM Tris-HCl and 270 mM glycine buffer (pH 8.4) for 16 h. All in-gel activity assays were carried out at 25 °C in the dark, with constant agitation. Gels were imaged using the ChemiDoc MP Imaging System, and band densities quantified with ImageJ.

### Metabolomics Analyses

GC-MS analyses were performed for samples of wandering L3 and mid-L2 larvae on an Agilent Technologies 5977B MSD–7890 GC system coupled with an autosampler, at the Mass Spectrometry Facility at Indiana University Bloomington, as previously described (Li and Tennessen, 2018). Identification and quantification of the metabolites shown in Figure 5 were performed using Mass Hunter Qualitative and Mass Hunter Quantitative software. The data was normalized by sample mass and an internal standard of succinic-d4 acid present at 2 μl/ml in the 90% methanol solution used for extraction.

Ultra-high-pressure LC-MS was conducted using a Vanquish UHPLC system coupled to a Q-Exactive mass spectrometer (Thermo Fisher Scientific, San Jose, CA, USA) at the University of Colorado Anschutz Medical Campus, following the configuration summarized in (Heidarian et al., 2023) and established protocols (Nemkov et al., 2019). The data was normalized by sample mass. MetaboAnalyst 6.0 (https://www.metaboanalyst.ca/) was used for metabolic pattern analysis through the unsupervised clustering method PCA or the supervised PLS-DA method, and for the construction of heatmaps. The data underwent log transformation (base 10) and Pareto scaling (Pang et al., 2021).

## Acknowledgements

The authors would like to thank Shefali Shefali and Dr. Carlos A. Couto-Lima for technical assistance. This work was supported by funding from the Fundação de Amparo à Pesquisa do Estado de São Paulo (FAPESP grants 2017/03806-7 and 2023/18123-3 to G.S.G.; 2022/01509-3 to M.F.O.; 2021/10910-0 to K.C.B.; and 2014/02253-6, 2021/06711-2, 2022/02370-9 and 2022/02371-5 to M.T.O.), the Conselho Nacional de Pesquisa e Desenvolvimento Científico (CNPq grants 141001/2019-4 to G.S.G.; and 306974/2017-7, 424562/2018-9 and 308940/2021-0 to M.T.O.), the Coordenação de Aperfeiçoamento de Pessoal de Nível Superior (CAPES fellowship to M.F.O; and CAPES-PrInt travel grant 88887.194785/2018-00 to J.L.), the National Institute of General Medical Sciences of the National Institutes of Health (MIRA grant 1R35GM119557 to J.M.T.), the Company of Biologists (Travelling Fellowship DEVTF2205766 to G.S.G.), and the American Society for Biochemistry and Molecular Biology (2022 PROLAB program to G.S.G.). INFABIC is supported by FAPESP (grant 2014/50938-8) and CNPq (grant 465699/2014-6).

## Supplemental Material

**Supplemental Figure Legends**

**Figure S1.**
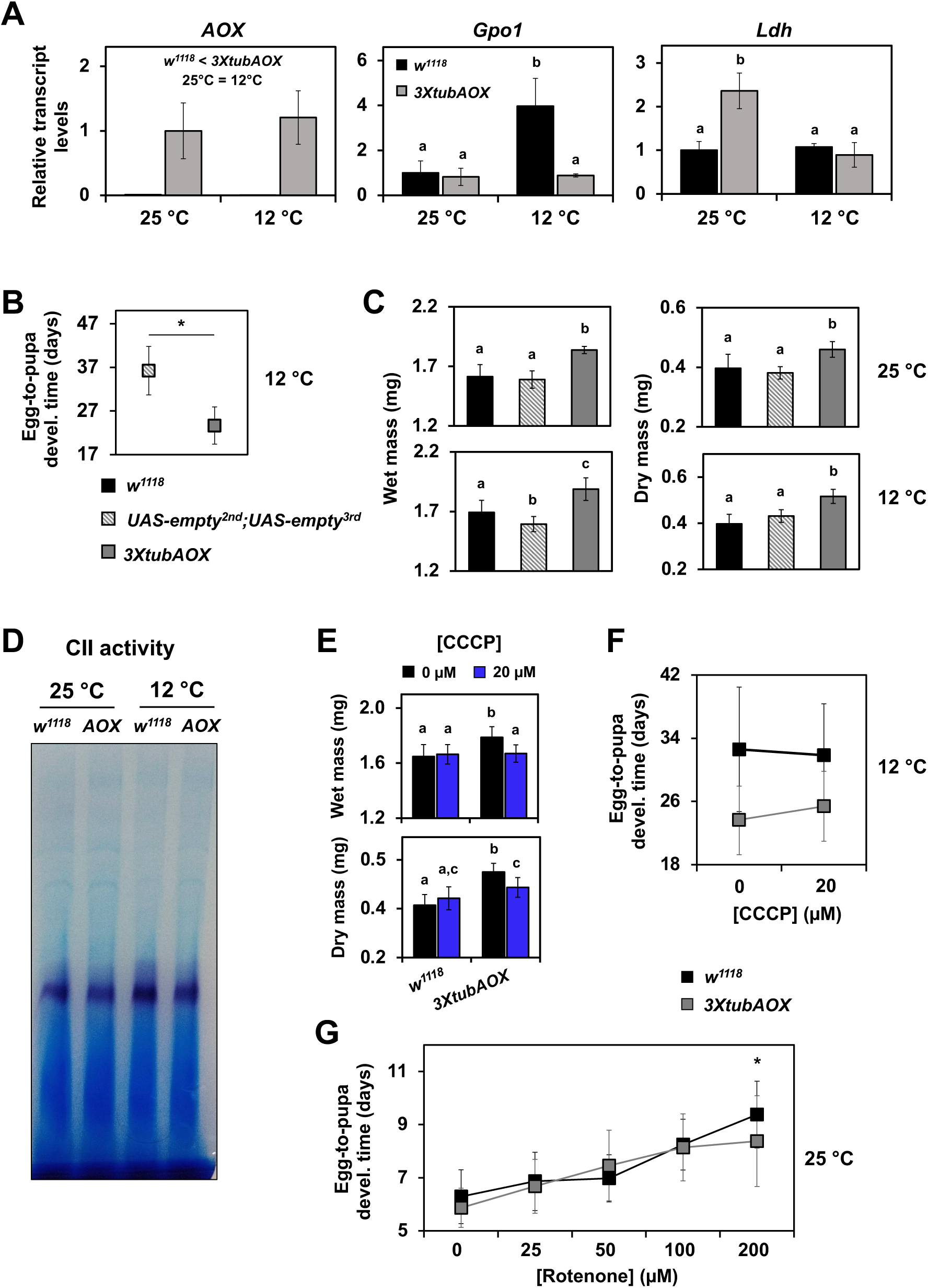
Effects of AOX expression on development. (A) Relative transcript levels of *AOX*, *Gpo1* and *Ldh* in wandering L3 larvae. Transcript levels were estimated using the levels of the housekeeping *eIF-1A* transcript as reference, and were normalized (arbitrarily set to 1.0) for *AOX* by its levels in AOX-expressing larvae (*3xtubAOX*) at 25 °C, and for *Gpo1* and *Ldh* by their levels in control larvae (*w^1118^*) also at 25 °C. The data represent the average of two independent biological replicates, and the error bars standard deviations. Egg-to-pupa developmental time (B) and larval body masses (C) at the indicated temperatures for the background control *w^1118^* line and the red-eye control *UAS-empty^2nd^;UAS-empty^3rd^* line, showing no effects of the *white* gene. (D) Representative results of in-gel activity for complex II (see Figure 3 for data quantitation). Larval body masses (E) and egg-to-pupa developmental time (F,G) of the *w^1118^* and *3XtubAOX* lines cultured on dietary supplementation with the uncoupler CCCP (E,F) and the CI inhibitor rotenone (G) at the indicated temperatures. Data represents the average of two-three biological replicates, each with at least five-ten technical replicates. * indicates significant differences according to Student’s t-test (p < 0.05). Two-way ANOVAs, followed by Tukey’s *post hoc* tests, were used to show differences (p < 0.05) between genotypes, between temperatures (denoted by the symbols < or =), or in the interaction genotype X temperature (denoted by the letters ‘a-c’).

**Figure S2.**
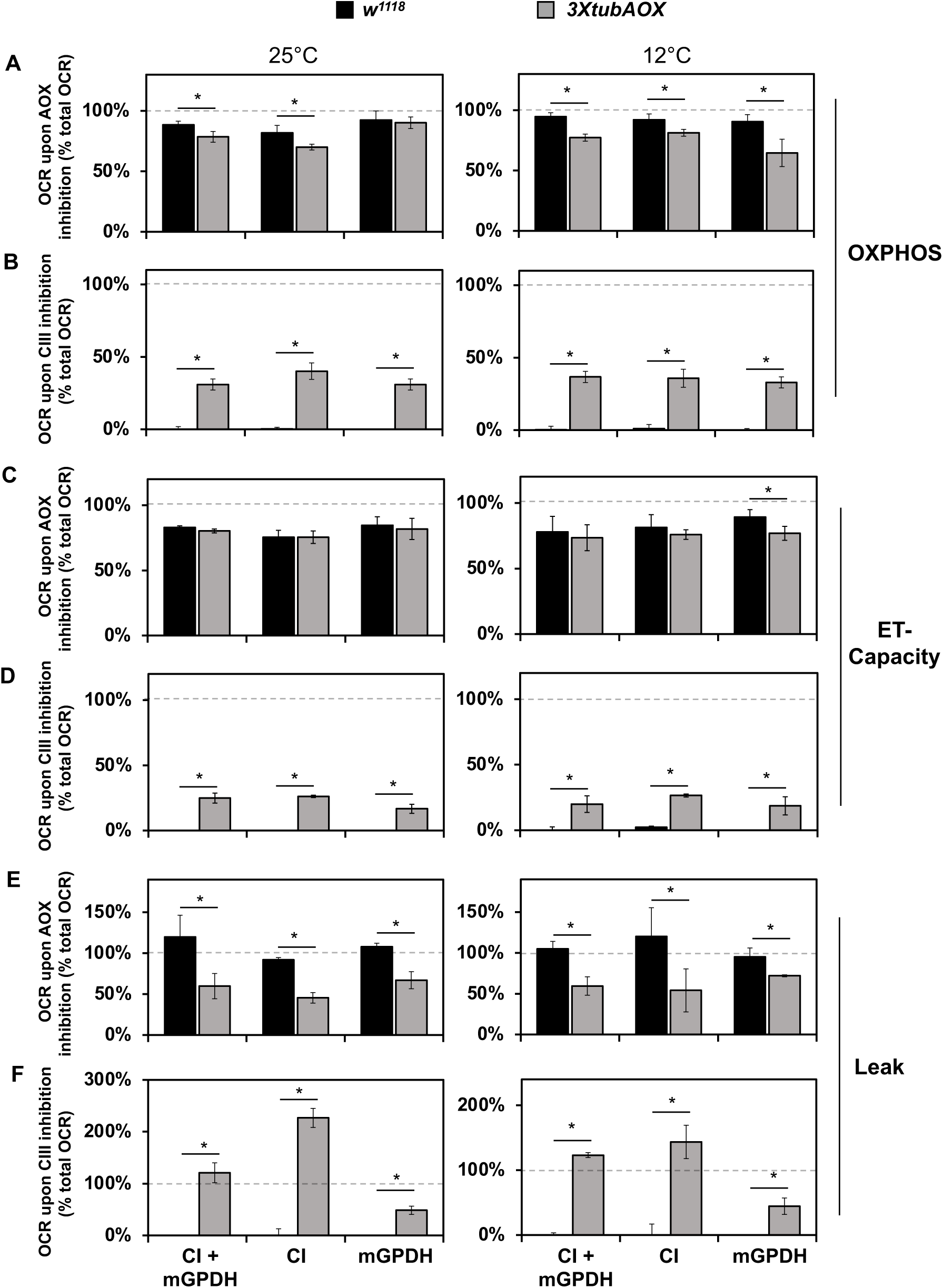
AOX significantly contributes to larval respiration, particularly to the Leak state. Remaining oxygen consumption rates (OCR) driven by the indicated dehydrogenase at the indicated temperature after inhibition of AOX with propyl gallate (A, C and E), or of CIII with antimycin A (B, D and F). The data represent percentages of the total OCR as shown in Figure 3A-C. The mitochondrial respiratory states OXPHOS (A and B), ET-capacity (C and D) and Leak (E and F) are shown. * indicates a significant difference (p < 0.05) between AOX and control larval samples, according to Student’s t-tests.

**Figure S3.**
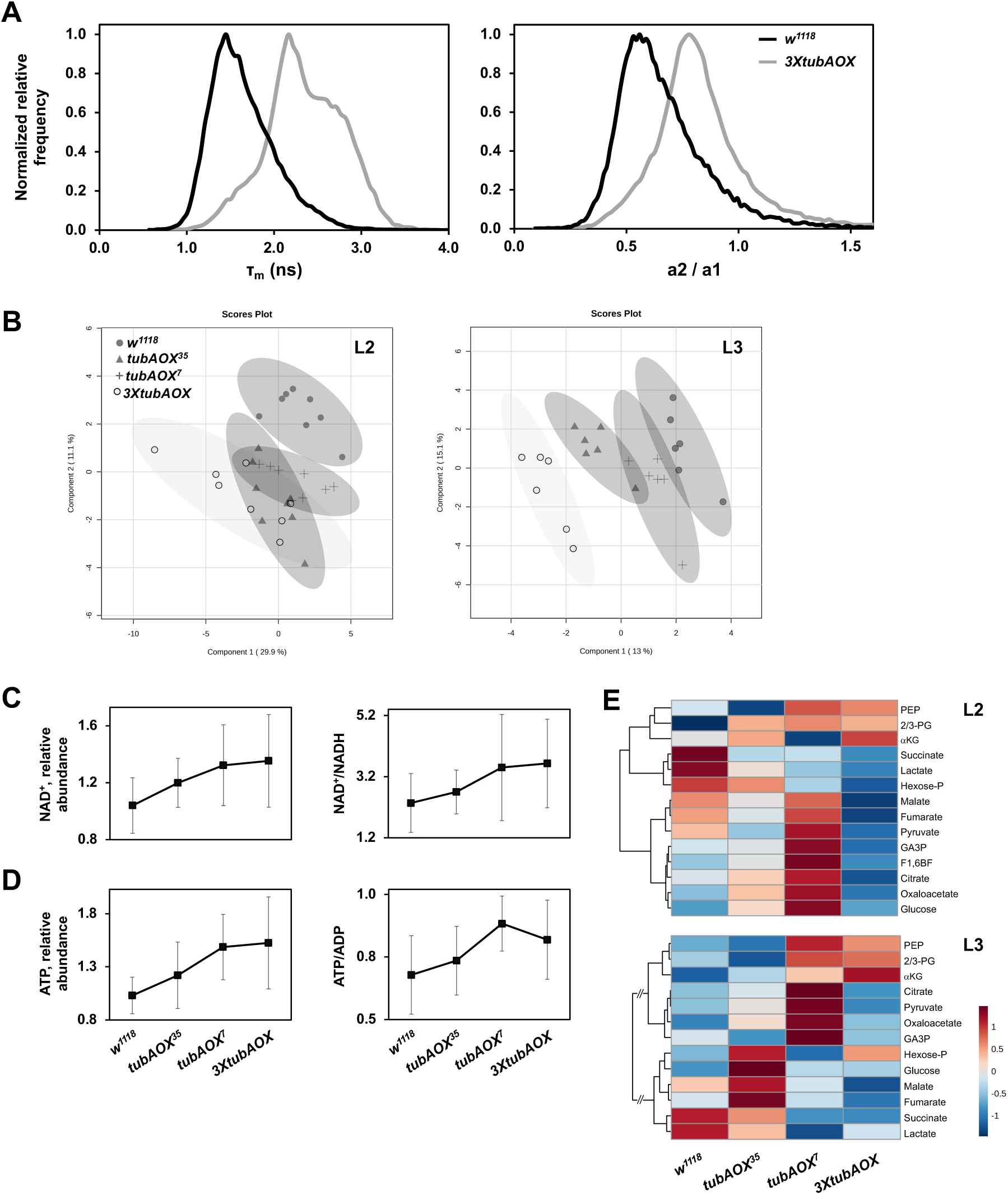
Metabolic analyses of AOX-expressing lines. (A) Distribution of τ_m_ of free NADH autofluorescence (left graph) and of reverse amplitude rate of enzyme-bound/free NADH (a2/a1, right graph) of samples of larval fat bodies of each genotype. (B) Partial Least Squares Discriminant Analysis (PLS-DA) plots showing separation among the metabolic profiles of mid-L2 and wandering L3 larvae of the indicated genotypes, based on liquid chromatography-mass spectrometry. (C,D) Tendency of AOX dose-dependent increase in NAD+ and ATP levels, and in the NAD+/NADH and ATP/ADP ratios. (E) Heatmaps indicating the relative levels of the identified glycolytic and tricarboxylic acid cycle metabolites.

## Supplemental Table

**Table S1:**
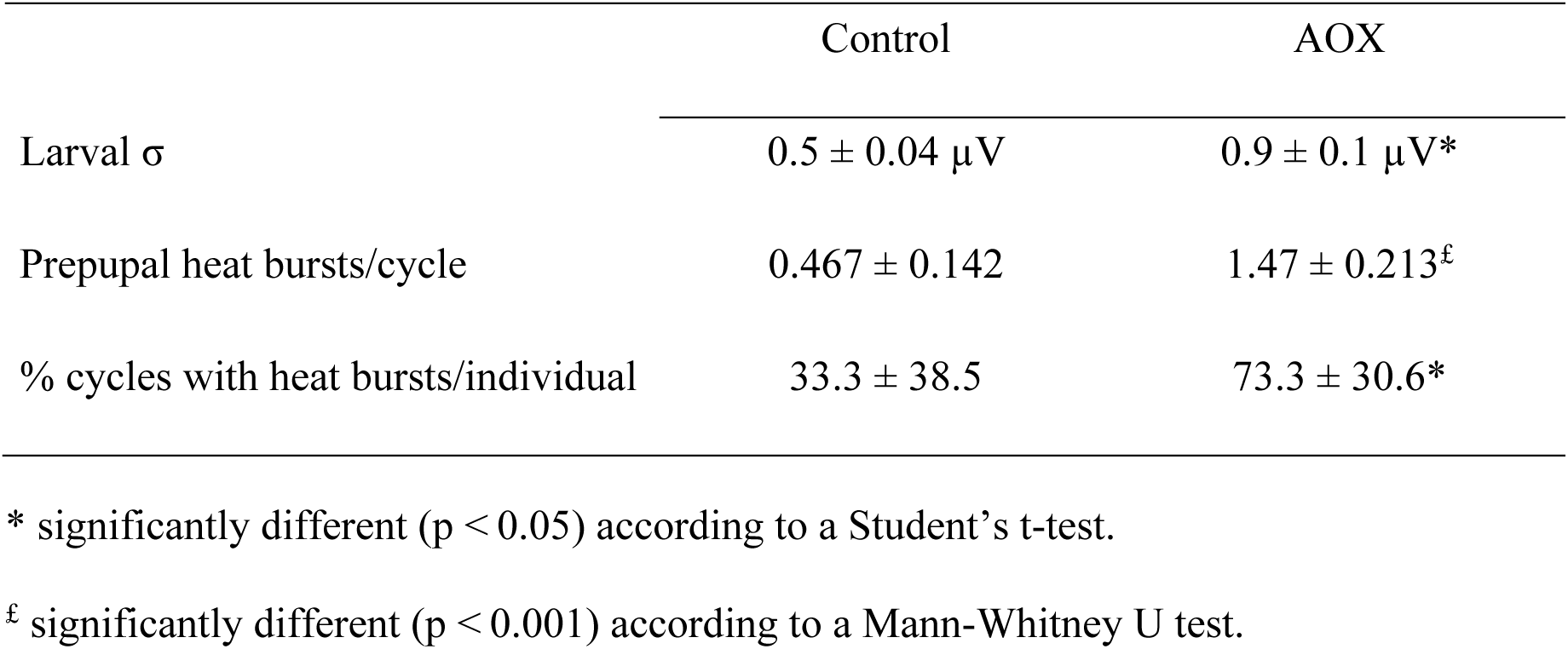
Larval agitation and prepupal heat burst events are increased by AOX. The average noise of the voltage change output of the chip calorimeter (σ, see representative output in Figure 1C) +/-standard error represents larval movements inside the calorimeter chambers. The data was obtained from two biological replicates, using 6-10 individuals in each one (see details in Material and Methods). The average number of prepupal heat bursts per cycle and the percentage of cycles with heat bursts per individual were calculated for 10 individuals from each genotype in the ∼18 min cycle of each real-time measurement of the calorimetry assays.

## Notes

### Competing Interest Statement

The authors have declared no competing interest.

